# A Midichloriaceae endosymbiont of terrestrial arthropods found as an endosymbiont in a marine nematode

**DOI:** 10.1101/2025.07.30.667539

**Authors:** Arno Hagenbeek, Yumiko Masukagami, Pradeep Palanichamy, Filip Husnik

## Abstract

The obligate endosymbiont *Candidatus* Lariskella (Alphaproteobacteria, *Candidatus* Midichloriaceae) has been found across a wide diversity of terrestrial arthropods, including ticks, true bugs, beetles, fleas, wasps and moths. However, to date, *Ca.* Lariskella had never been detected in nematodes or marine animals. Here we report the first known occurrence of *Ca*. Lariskella infecting a population of marine nematodes (Enoplida, Thoracostomopsideae). This nematode-infecting *Ca.* Lariskella is closely related to insect-infecting *Ca.* Lariskella, despite the drastic shift in both host phylum and habitat. TEM and FISH microscopy showed *Ca.* Lariskella is localized within both somatic cells and developing oocytes, strongly suggesting that *Ca.* Lariskella is a vertically transmitted endosymbiont within the nematode population. We found that *Ca.* Lariskella was present within approximately 20% of the nematode population, but notably failed to detect any *Ca.* Lariskella within adult males, potentially hinting at reproductive manipulation. Overall, our findings show that *Ca*. Lariskella is not limited to arthropods or terrestrial hosts, indicating a larger host range than previously described. Its presence within marine nematodes demonstrates the ability of *Ca.* Lariskella to infect nematode hosts, as well as hosts from marine environments, suggesting terrestrial nematodes, marine arthropods, and perhaps even other marine invertebrates could be potential hosts of *Ca.* Lariskella.

## Introduction

The Rickettsiales are a large, well-studied order within Alphaproteobacteria, best known for being obligate endosymbionts. Due to their endosymbiotic nature, Rickettsiales typically possess small genomes (1-1.5Mb) with streamlined gene contents [1]. Rickettsiales have been widely studied for decades, initially primarily for their role in diseases such as typhus (*Rickettsia prowazekii*), Rocky Mountain spotted fever (*Rickettsia rickettsii*), and anaplasmosis (*Anaplasma phagocytophilum*) [2]. However, diverse Rickettsiales have also garnered increasing evolutionary interest, as their obligate symbiotic lifestyle resulted in a wide array of unique adaptations, such as supplementing host nutrition [3], aiding host defense [4], manipulating host apoptosis [5], or manipulating host sex determination [6].

*Candidatus* Midichloriaceae (hereafter Midichloriaceae) comprises a relatively novel family within the Rickettsiales order, having been formally described only in 2013 [7]. Currently, Midichloriaceae represent by far the least studied among the three families of Rickettsiales, in part due to their novelty, and in part because many Midichloriaceae infect poorly sampled host taxa such as protists [8]. Despite being less studied, Midichloriaceae have been discovered across a vast phylogenetic range of hosts such as in multiple protist supergroups [9–12], placozoans [13,14], corals [15], multiple arthropods [16– 21], as well as in secondary infections of mammals [22–24]. Additionally, Midichloriaceae have been found to exhibit several unique evolutionary traits, such as being the only group in Rickettsiales known to maintain flagellar genes [8,25,26], and including the only known bacterial species that can reside inside the host mitochondria [21].

Midichloriaceae appear to primarily infect aquatic hosts. However, the family contains two ‘terrestrial’ genera: *Candidatus* Lariskella (hereafter *Lariskella)* and *Candidatus* Midichloria (hereafter *Midichloria)*. Both *Lariskella* and *Midichloria* have primarily been studied in ticks, where they appear to be widespread [16,21,27,28] and are hypothesized to play a nutritional role [29]. Notably, both *Lariskella* and *Midichloria* are transmitted to mammals during the tick’s blood meal, though it has not been confirmed whether either bacteria can replicate within mammals or cause disease [22–24].

While *Midichloria* is only known to infect ticks, *Lariskella* infects a far broader range of arthropod hosts. It has been reported in ticks [16,27,28,30], stinkbugs [19], leaf-footed bugs [31], weevils [17], psyllids [18], beetles [20], and fleas [32]. In addition to these published reports, *Lariskella* genomes have been sequenced from wasps (NCBI Refseq GCF_964020185.1) and moths (NCBI Refseq GCF_964019805.1). Here we add to this host range, reporting the occurrence of *Lariskella* as an endosymbiont of a marine nematode (Enoplida, Thoracostomopsideae). This nematode represents the first non-arthropod host of *Lariskella*, as well as the first known occurence of *Lariskella* infecting an aquatic host. We assembled a metagenome-assembled genome (MAG) of the nematode-infecting *Lariskella.* This MAG showed high genetic similarity to insect-infecting *Lariskella* despite the drastic host shift. Functional annotation showed that *Lariskella* is capable of synthesizing several essential B vitamins. Using microscopy we demonstrate that the nematode-infecting *Lariskella* is an intracellular endosymbiont of the nematode species, and is likely maternally transmitted. Lastly, we show that *Lariskella* is present at low frequencies (20%) in the nematode population, and appears to be completely absent from males. This infection pattern could indicate reproductive manipulation, although we could not confirm this experimentally due to the inability to culture the nematode.

## Materials and methods

### Nematode sampling

From March 2023 until June 2024, sediment samples were taken from the Okinawan coast (26.4723631548844, 127.83168712213792). Nematodes were extracted from the sediment through a magnesium chloride decanter method: 100 mL of 7.5% MgCl_2_ was added to approximately 200 mL of sediment. Sediment was stirred with a spoon and left to sit for 10 min, after which it was stirred again, and the solution was sieved through a 32 μm sieve. The contents of the sieve were washed into a Petri dish using sand-filtered seawater. Nematodes were then picked from the Petri dish using an Irwin loop under a dissecting microscope (Olympus SZX16). Nematodes picked for microscopy were washed once in filter-sterilized (0.22 μm) seawater to remove debris and then transferred directly to the appropriate fixative. All other nematodes were cleared of surface contaminants by a triple serial transfer in filter-sterilized seawater using sterile pipette tips. Cleaned nematodes were then frozen at −70°C awaiting processing.

### Illumina sequencing

For 15 nematodes, shotgun metagenomes were obtained using Illumina sequencing. DNA was extracted from the nematodes using the MasterPure Complete DNA and RNA purification kit (Thermo Fisher Scientific). The kit was used following the manufacturer’s instructions, with the exception of a prolonged (2 h) proteinase K incubation to facilitate full lysis of the nematode. Illumina libraries were prepared using the NEBNext Ultra II PCR plus kit (New England Biolabs) and sequenced on the SP flow cell of the NovaSeq 6000 platform with 150 bp paired-end reads. The generated reads were thereafter quality checked using Fastqc (v0.11.9) (http://www.bioinformatics.babraham.ac.uk/projects/fastqc/).

### Host 18S rRNA gene sequencing and phylogeny

The 18S rRNA gene sequences of the nematode host were assembled from the metagenomes and approximately identified using phyloFlash (v3.4) [33]. To determine the host’s exact phylogenetic position, an 18S rRNA gene tree was reconstructed. Based on the phyloFlash results, the nematode was related to the order Enoplida. Therefore, 18S rRNA gene sequences of nematodes from Enoplida were downloaded from SILVA [34]. Triplonchida 18S rRNA gene sequences were used as an outgroup. The downloaded sequences were aligned with the metagenome-reconstructed 18S rRNA gene sequence using MAFFT (v7.505) [35] set to a maximum of 1,000 iterations, and using the -localpair argument. A phylogenetic tree was reconstructed from the alignment using IQ-TREE2 (v2.0.7) [36] set to 1,000 bootstraps using the GTR+F+R4 model selected by the IQ-TREE2 modelfinder based on the Bayesian Information Criterion. As a comparison, trees were also reconstructed using the GTR+G+I model frequently used in nematode 18S rRNA gene phylogeny literature [37,38] and both trees had identical topologies with only minor differences in branch lengths (Supplementary figure S1). The tree suggested that the *Lariskella* host was related to *Mesacanthoides,* and we confirmed the 18S rRNA gene sequence similarity using BLASTn megablast on the NCBI website (3-6-2025) [39].

While the Illumina results suggested the nematode population at the sampling site was uniformly one species, we additionally confirmed the species identity by Sanger sequencing the 18S rRNA gene from 10 individuals, randomly selected from several sediment samples. The PCR was performed with the PF1 [40] and FAD4 [41] eukaryote 18S rRNA gene primers, using the Q5 high-fidelity DNA polymerase (New England Biolabs). Thermocycling conditions were as follows: initial denaturation at 98**°**C for 30 s; then 30 cycles of 98°C for 10 s, 65°C for 25 s, 72°C for 60 s; followed by a final elongation at 72°C for 2 min. Sanger sequencing was performed by first cleaning the PCR products using the Monarch PCR & DNA Cleanup Kit (New England Biolabs). Fluorescent labeling was then done using the BigDye Terminator v3.1 cycle sequencing kit (ThermoFisher Scientific) according to the manufacturer’s protocol, followed by cleaning using Agencourt CleanSEQ (Beckman Coulter). The cleaned products were eluted into DNase- and RNase-free water and sequenced by a SeqStudio Genetic Analyzer (ThermoFisher Scientific).

Obtained sequences were compared against the nematode 18S rRNA gene reconstructed from the Illumina sequencing metagenome to confirm the species. As Sanger sequencing further confirmed the uniformity of the population, it was thereafter only confirmed through examination of nematode morphology.

### *Lariskella* identity and diagnostic PCR

The initial identification of *Lariskella* was performed by 16S rRNA gene reconstruction using phyloFlash. To experimentally confirm the presence of *Lariskella*, a diagnostic PCR with *Lariskella*-specific primers was performed in 10 nematodes. We amplified the *Lariskella* rrl-rrf intergenic spacer using the *Lariskella-*specific Mtz-5S-F and Mtz-23S-R primers [42] and Q5 high-fidelity DNA polymerase (New England Biolabs), with the following thermocycling conditions: initial denaturation at 94**°**C for 4 min; then 30 cycles of 94°C for 1 min, 62°C for 1 min, 72°C for 80 s; followed by a final elongation at 72°C for 4 min. To confirm that the bands were the intended products, the obtained bands were Sanger sequenced as described above. Resulting Sanger sequences were compared against the *Lariskella* rRNA operon obtained during Illumina sequencing using EMBOSS Needle [43], to ensure the PCR product originated from the *Lariskella* symbiont.

### *Lariskella* genome assembly and phylogeny

Based on the 16S and 18S rRNA gene sequences, both the host (N=15) and *Lariskella* (N=4) were identical across the 15 shotgun-sequenced nematode samples. The reads for these samples were thereafter trimmed using Fastp (v0.23.2) [44] and co-assembled using Megahit (v1.2.9) [45]. The coverage for this assembly was then calculated by Bowtie2 (v2.4.2) [46], and sorted, compressed, and indexed using SAMtools (v1.12) [47]. The assembly was then binned using Metabat2 (v2.12.1) [48], CONCOCT (v1.1.0) [49], and SemiBin2 (v2.0.2) [50], and the resulting bins were refined with DASTool (v1.1.6) [51]. It was then assessed which of the metagenome assembled genomes (MAGs) corresponded to *Lariskella* through a threefold method: first, a MEGAN (v6.23.3) [52] lowest common ancestor assignment analysis was run to confirm the approximate composition of the MAG. Second, the 16S rRNA genes in the MAGs were predicted using barrnap (v0.9) (https://github.com/tseemann/barrnap), and aligned by BLASTn (v2.10.0) [39] against the 16S rRNA gene reconstructed by phyloFlash. Finally, the MAG was aligned with the Mummer nucmer algorithm (v3.23) [53] against the complete *Lariskella* reference genomes obtained from NCBI (GCF_964019805.1 and GCF_964020185.1). For all MAGs of *Lariskella* origin, the best MAG was chosen based on completeness and duplication scores inferred by BUSCO (v5.4.4) [54] using the Rickettsiales_odb10 database. The best quality MAG was found to originate from CONCOCT. CONCOCT filters out contigs smaller than 1000bp by default, and these are thus not present within the *Lariskella* MAG.

The phylogenetic position of the new *Lariskella* MAG was inferred from a multigene alignment of Midichloriaceae. Midichloriaceae reference sequences were downloaded from NCBI, with Anaplasmataceae and Rickettsiaceae sequences selected as outgroups. Prokka (v1.14.6) [55] was used to predict protein-coding sequences, which were piped into Orthofinder (v2.5.4) [56] to find singlecopy orthologs shared across the 25 genomes. The protein sequences of the 141 shared single-copy orthologs were aligned using MAFFT (set to -localpair, with 1,000 iterations), trimmed using trimAl (v1.4.rev22) [57] set to automated1, and concatenated with a python script. A maximum likelihood phylogenetic tree was reconstructed from the concatenated alignment using IQ-TREE2, set to 1,000 bootstraps. IQ-TREE2 model finder chose LG+F+R5 as the best model based on the Bayesian Information Criterion. As a comparison, trees were reconstructed using the LG+F+I+G4 and LG+C60+I+G4 models used in Giannotti et al. (2022) [8], both of which had similar topologies to the LG+F+R5 tree (Supplementary figure S2).

### Nutritional contribution analysis

To infer any nutritional contribution by *Lariskella*, the completion of metabolic pathways synthesizing nutrients essential for nematodes was assessed using the genome data. Essential nutrients were based on findings in *Caenorhabditis elegans* [58]. The two reference *Lariskella* genomes and the nematode *Lariskella* MAG were annotated using EggNOG (v2.1.12) [59] in genome mode, using the bundled MMseqs2 BLASTx as the gene prediction method, and the EggNOG bacterial database. The KEGG KO numbers were extracted from the .annotations EggNOG output file using a Python script, and used to assess the completion of the KEGG modules corresponding to the synthesis of essential nutrients in KEGG mapper reconstruct [60,61]. If there were multiple redundant KEGG modules, only the most complete bacterial pathway was considered.

### DOPE-FISH microscopy

Multiple nematodes (5-20 individuals per slide) were picked and placed in a 1 mL drop of filter-sterilized (0.22 μm) sea water. They were then transferred to a 1.5 mL tube containing 100 μl of 1:1 glacial acetic acid:ethanol solution, where they were fixed for 10 min. Fixed nematodes were washed in 100% ethanol two times for 5 min, and incubated in 1:1 methanol:PBS-T (1X PBS buffer containing 0.1% Tween 20) for 10 min. Afterwards, they were incubated in 1% paraformaldehyde (Nacalai) PBS-T, which was washed away with two PBS-T washes of 2 min each. Nematodes were prepared for hybridization with three washes in hybridization buffer (0.9 M NaCl, 0.02 M Tris-HCl, 0.01% sodium dodecyl sulphate (SDS, Sigma-Aldrich) + 30% formamide; Wako) of 5 min each. The nematodes were then hybridized in 100 μL of hybridization buffer containing DAPI DNA stain (0.01 μg/μL; Roche), and 0.1 μM of *Lariskella*-specific probe (JBioS) overnight at 42°C. The *Lariskella*-specific probe (CTGTGATAGTCCGGCCGAAC) was purified through high performance liquid chromatography purification. This probe was then labelled at both the 5’ and 3’ ends with Alexa Fluor 647. The DNA sequence of the probe was determined to be fully *Lariskella*-specific using SILVA’s TestProbe (v3.0) function [34]. After hybridization, excess DAPI and probe were washed away with three PBST washes of 10 min. Nematodes were suspended in MilliQ water and pipetted onto a glass slide. After removing excess water, the nematodes were mounted using ProLong Diamond Antifade Mountant (Thermo Fisher Scientific) and imaged using a confocal fluorescence microscope (Microscope: Nikon A1, detector:x2 MA PMT, x2 GaAsp, x1 T-PMT, lasers: 404.6 nm and 637.8 nm).

### Transmission electron microscopy (TEM)

Five adult female nematodes were picked from sediment samples. These nematodes were fixed by briefly (2 min) submerging in the fixative (2.5% glutaraldehyde; Fujifilm Wako and 2% paraformaldehyde; Fujifilm Wako in 0.1 M HEPES; Nacalai). The anterior of the nematode (from the mouth to slightly behind the end of the pharynx) was dissected from each nematode. The posterior of the nematodes was incubated in the fixing solution overnight at 4°C, awaiting TEM post-fixation steps. The processed nematodes were assessed for *Lariskella* infection by performing the *Lariskella* diagnostic PCR described above on the dissected pharynges. One nematode was found to be *Lariskella*-positive, and its posterior was prepared for TEM. The posterior of the *Lariskella-*positive nematode was washed with 0.1 M HEPES for 5 minutes three times, and post-fixed with 2% osmium tetroxide for 1 hour at room temperature. After post-fixation, the nematode was *en bloc* stained with 2% uranyl acetate overnight at 4°C. The nematode was dehydrated by serial transfer through increasing concentrations of ethanol (30%, 50%, 70%, 90%, 100%), incubating for 10 minutes in each, and two times 10 minutes in 100% ethanol. This was followed by a final dehydration using 100% acetone for two times 10 minutes. Low viscosity resin (Agar Scientific, catalog no. R1078) was then infiltrated by replacing with 2:1 (vol:vol) of acetone:resin for 2 hours at room temperature while rotating in an TAITEC Rotator RT-50. After 100% of the resin was replaced, the resin was infiltrated under a 1-hour vacuum and incubated overnight at room temperature. The resin was then polymerized for 48 hours at 60°C. The resin block was trimmed using glass knives, and ultrathin sections (50-60 nm) were cut using a Leica Ultramicrotome EM (UC7). Ultrathin sections were collected onto the 0.5% formvar (SPI-CHEM) coated copper one-slot grids (Nisshin-EM). The air-dried grids were then examined using a JEOL 1400 Flash transmission electron microscope.

### Sex-specific infection rate experiment

*Lariskella* diagnostic PCR was performed on adult nematodes of known sex to determine per-sex infection rates. Nematodes were collected, and their sex was determined based on morphology using an inverted microscope (Nikon Eclipse Ti2-E). Due to the difficulty of determining sex in juveniles, only adults were considered. Adults with a sclerotized gubernaculum were determined as males. The gubernacula had clear autofluorescence under 488 nm light, which was used to more easily identify adult males. Adults lacking a gubernaculum, but containing ovaries with visible eggs, were considered as fully-developed adult females. No hermaphrodites were observed. The presence of *Lariskella* was then confirmed in the sexed nematodes using the *Lariskella* diagnostic PCR described above. In total, 35 sexed nematodes were tested (23 females, 12 males).

### Analysis of *wmk* ortholog presence in *Lariskella*

To identify potential male-killing genes in the nematode *Lariskella*, we compiled a curated reference set of *Wolbachia*-derived WO-mediated killing (*wmk*) genes. The *Lariskella* genomes were screened for orthologs using OrthoFinder. To improve the resolution of this analysis, we generated custom BLAST databases containing the reference *wmk* gene sequences. Protein-coding genes from *Lariskella* were predicted using Prokka and queried against this database using BLASTp (E-value ≤ 1e^−5^, minimum sequence identity ≥ 30%).

The protein domains present in the inferred *wmk* orthologs were identified using InterProScan (v5.60-92.0) [62]. InterProScan reported only 1 domain shared among all *wmk* orthologs, which was annotated as a Cro/C1-type DNA-binding domain. Alphafold2 (v2.1.1) [63] was used to infer the protein structures, which were visualized in iCn3D [64]. Structural alignments were performed using TM-align. These structural alignments showed a helix-turn-helix structure shared among all identified orthologs, the position of which corresponded to the Cro/C1-type DNA-binding domains reported by InterProScan.

## Results

### Nematode host phylogeny

To infer the phylogeny of the nematode host, its 18S rRNA gene sequence was obtained by both 18S rRNA Sanger sequencing and metagenome assembly. The 18S rRNA gene sequence was used to reconstruct a host phylogenetic tree (Fig. 1). The host 18S rRNA gene did not correspond to any of the available nematode 18S rRNA sequences but was placed within the family Thoracostomopsidae of the order Enoplida. The nematode host clusters together with a *Mesacanthoides* species, although this assignment is uncertain, as only a single *Mesacanthoides* 18S rRNA gene was available. The 16 rRNA gene of the *Lariskella-*infected nematode shared 97% sequence identity with this *Mesacanthoides* sequence.

**Figure 1:**
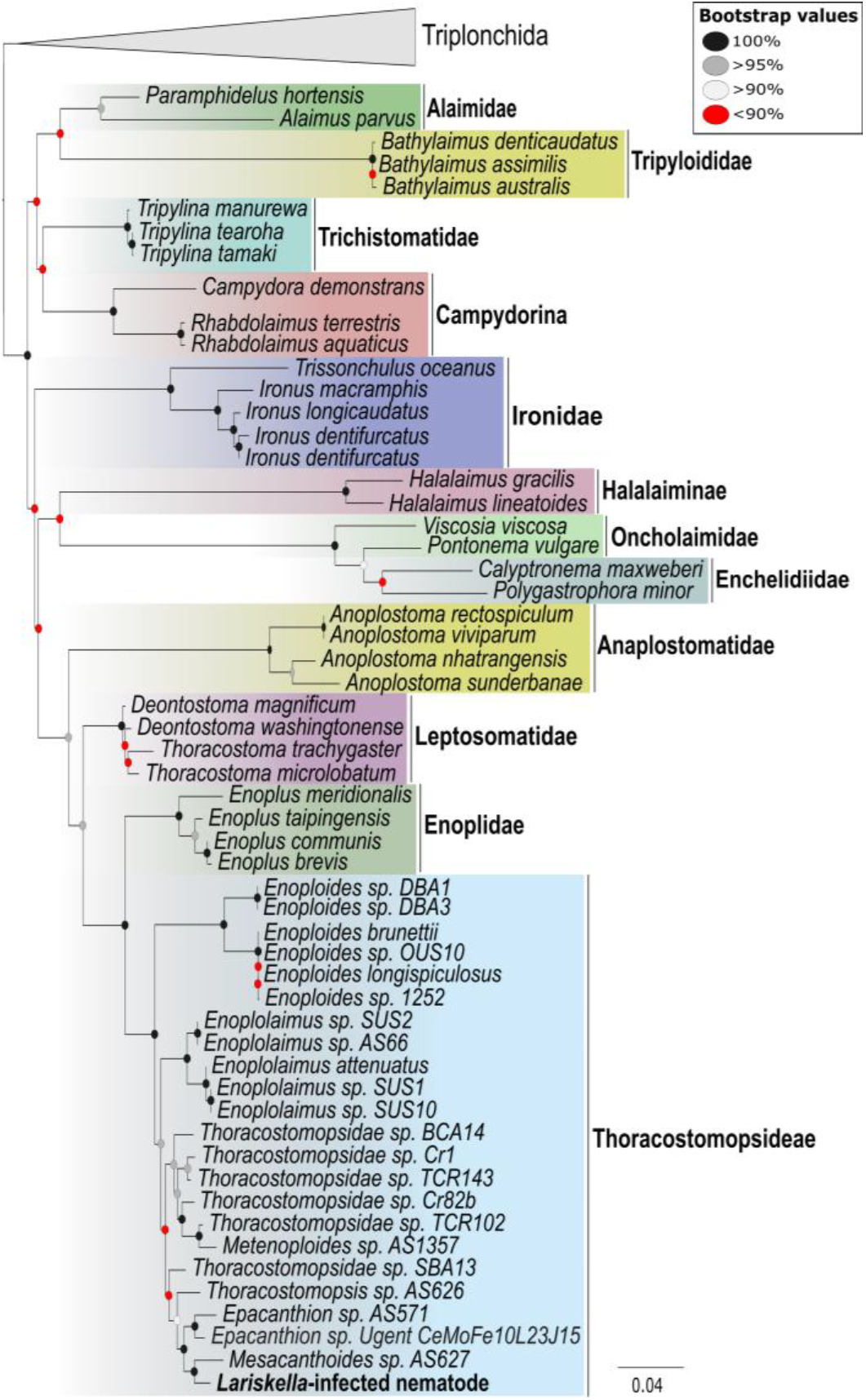
Phylogenetic tree of the nematode host. Phylogenetic tree based on nematode 18S rRNA gene sequence, demonstrating that the nematode host belongs to the Thoracostomposidae. The tree was inferred by maximum-likelihood based on the GTR+F+R4 model.

### *Lariskella* identity, genome, and phylogeny

The *Lariskella* infection in nematodes was first discovered by the presence of a *Lariskella arthropodarum-*related (99.6% identity) 16S rRNA gene in the nematode metagenome. Diagnostic PCRs with *Lariskella*-specific intergenic primers were performed to experimentally confirm the presence of *Lariskella*. In total (including the sex-specific infection rate experiment), diagnostic PCRs were performed on 50 individuals, of which ten gave visible bands indicating *Lariskella* infection.

Using binning, a MAG of the suspected *Lariskella* was obtained from 15 co-assembled nematode metagenomes. This MAG was compared against two publicly available *Lariskella* genomes. The MAG is fragmented (321 contigs) but is highly similar to the references in assembly size (1.5Mb), BUSCO completeness score (93%), and GC content (35%) (Supplementary Table S1). The MAG was aligned against the references and showed a high degree of nucleotide-level similarity, further confirming its identity as *Lariskella* (Supplementary figure S3).

To assess the phylogenetic position of this novel *Lariskella* lineage, a 141-gene multigene phylogenetic tree was reconstructed for the *Midichloriaceae* family (Fig. 2).

**Figure 2:**
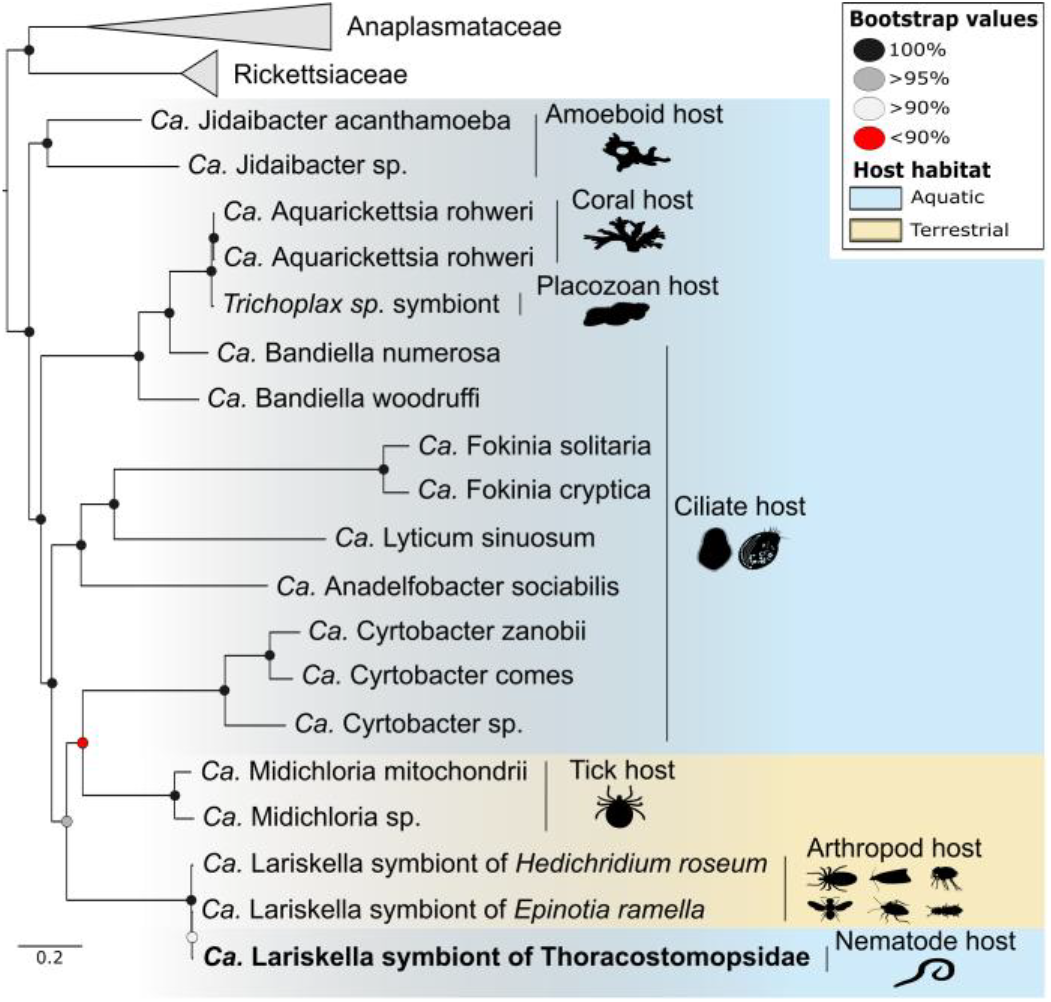
Phylogenetic tree of Midichloriaceae. 141-gene multigene phylogenetic tree of the family Midichloriaceae. The tree shows that the nematode-infecting *Lariskella* clusters among the insect-infecting *Lariskella* with minimal genetic divergence. The tree was inferred by maximum-likelihood based on the LG+F+R5 model.

The tree clearly places the *Lariskella* MAG among the *Lariskella* reference genomes. Notably, despite colonizing a host from a completely different phylum, the nematode-infecting *Lariskella* placed among the insect-infecting *Lariskella* with negligible branch lengths.

### *Lariskella* metabolic capacity

The *Lariskella* genomes encode complete pathways required to synthesize the essential vitamins pyridoxine (B6) and biotin (B7) (Fig 3). However, biosynthesis pathways of none of the essential amino acids were fully encoded by any of the *Lariskella* genomes. The pathways for lysine, heme, and tetrahydrofolate are nearly complete, and could potentially be completed by uncharacterized enzymes or through substrate exchange with the host.

**Figure 3:**
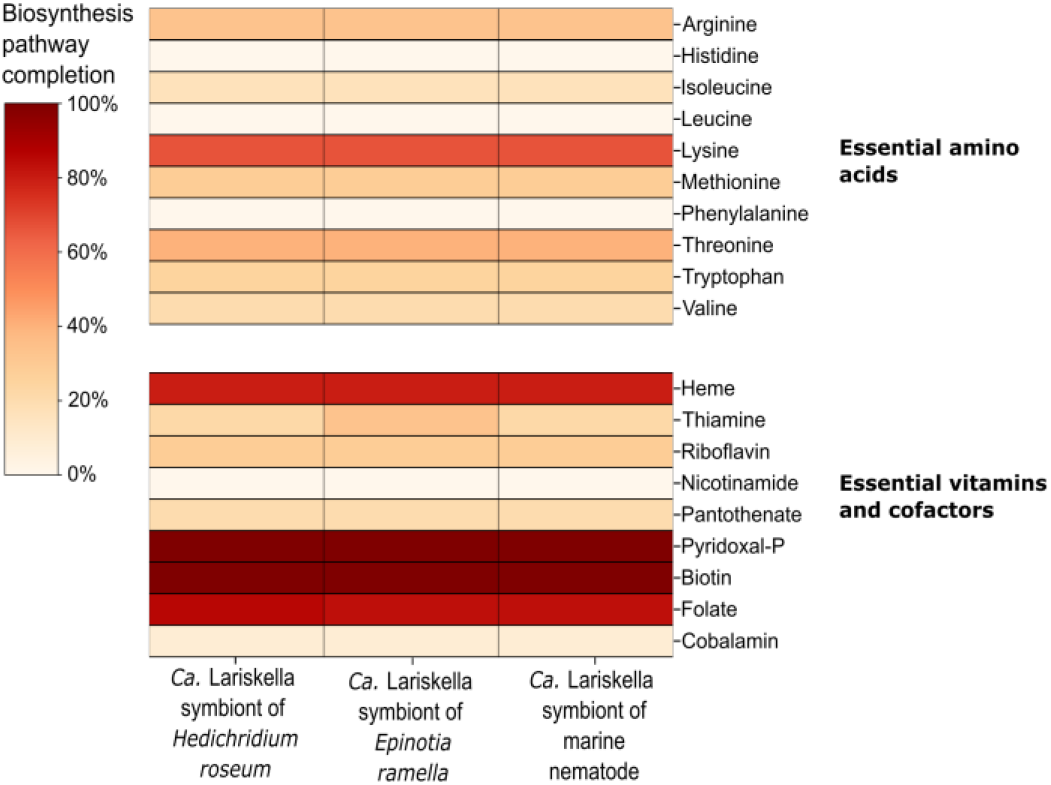
Metabolic capacity of *Lariskella*. Heatmap displaying the completion of biosynthesis pathways in the *Lariskella* genomes. Pathway completion was found by calculating KEGG module completion based on EggNOG K-term output. The shown pathways correspond to nutrients essential to nematodes, based on findings in *Caenorhabditis elegans*.

### DOPE-FISH microscopy and TEM

*Lariskella* localization was visualized using DOPE-FISH microscopy (Fig. 4A-C). The localization of the *Lariskella*specific probe shows *Lariskella* has a patchy distribution of intracellular infections throughout the somatic tissue of the nematode host. The ovaries of the nematode clearly deviate from this patchy distribution, and instead display highly abundant infections of *Lariskella* in both the maternal ovarian tissue and the developing oocytes.

**Figure 4:**
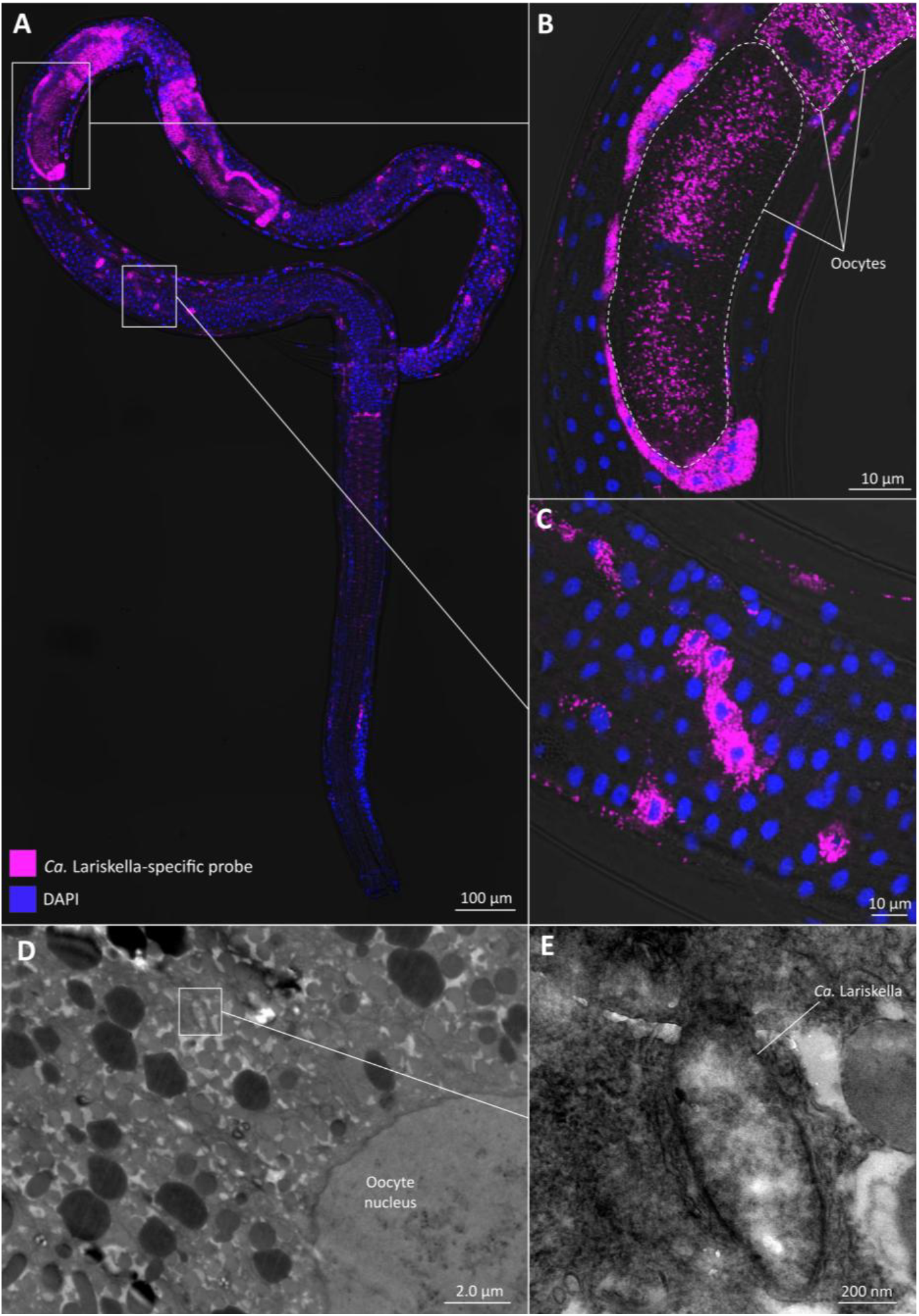
Microscopy of the nematode-infecting *Lariskella*. A to C) DOPE-FISH microscopy images of a whole-mounted gravid female. Pink indicates the *Lariskella*-specific probe. Blue indicates DAPI, which stains all DNA and visualizes host nuclei. A) Whole nematode. B) Zoomed image of one of the paired ovaries, including infected oocytes C) Zoomed image of somatic tissue, including gut epithelium and cuticle. D and E) TEM micrographs of a nematode ovary. D) Electron micrograph showing the inside of an infected oocyte. E) Electron micrograph within an oocyte, zoomed in on a *Lariskella* cell.

To further confirm the intracellular nature of the nematode *Lariskella*, TEM micrographs of various nematode tissues were inspected. The micrographs show the presence of bacteria-like structures within the oocytes (Fig. 4D-E), as well as within the somatic cells (Supplementary figure S4).

### Sex-specific infection rates

Diagnostic PCR showed an infection rate of 26% for female nematodes (6/23 infected) (Fig 5). Adult males were found to be free of infection, with none of the tested males (N=12) showing bands during the PCR. These results matched the DOPE-FISH results, where no *Lariskella* signal was observed in any of the processed males.

**Figure 5:**
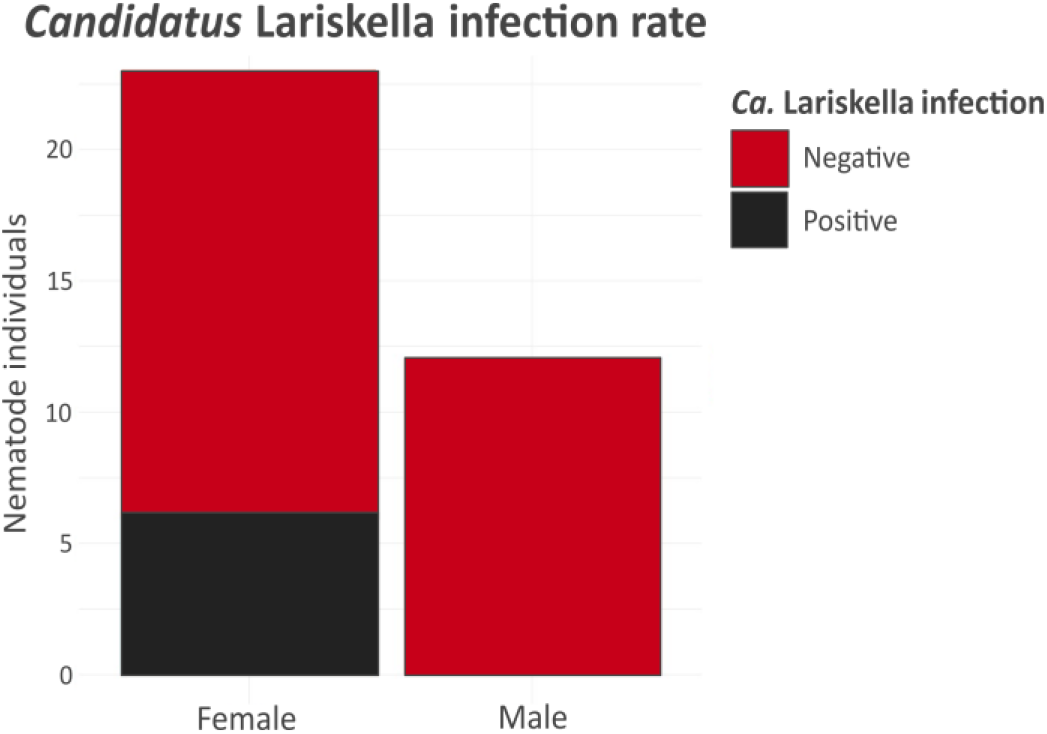
Infection rate of *Lariskella* within nematodes. Bar plot displaying the infection status of tested nematodes. Sexed nematodes were examined for *Lariskella* infection using diagnostic PCR (N=35).

## Discussion

In this study, a wild population of marine nematodes was confirmed to be infected by *Lariskella* through multiple experiments. The presence and identity of the *Lariskella* symbiont were confirmed through metagenomics, diagnostic PCR, and DOPE-FISH microscopy. In addition, the intracellular localization of the nematode-infecting *Lariskella* was confirmed using DOPE-FISH microscopy and TEM. All results point towards the *Lariskella* being a vertically-transmitted endosymbiont of the nematode host. This novel host represents the first known nematode host of the genus *Lariskella*, as well as the first known nematode host within the family *Midichloriaceae*. Additionally, this marine nematode represents the first known aquatic host found for *Lariskella,* which was previously considered exclusively terrestrial [8].

### Expanded host range of *Lariskella*

*Lariskella* was already known to have a broad host range, which this study expands by including nematodes. Curiously, with this novel addition of nematodes, the *Lariskella* host range roughly resembles the host range of *Wolbachia* (Supplementary figure S5). *Wolbachia* is a group of highly studied Alphaproteobacterial symbionts famous for their success as endosymbionts, being estimated to infect 50% of all arthropods [65]. *Wolbachia* are known to infect arachnids and insects [65], as well as nematodes [66,67], all of which are now known to also harbor *Lariskella*. Notably, *Wolbachia* also infect crustaceans [68], for which no *Lariskella* have currently been discovered. We suspect further sampling of crustaceans would reveal *Lariskella* infections, particularly as our findings illustrate the ability of *Lariskella* to infect marine hosts. If this is indeed true, it would raise interesting questions regarding Rickettsiales evolution and host choice, as *Wolbachia* and *Lariskella* are only loosely related, and yet seem to have independently converged on a similar host range.

### The phylogeny suggests a recent infection of nematodes

It was unexpected for the nematode-infecting *Lariskella* to be so closely related to the insect-infecting *Lariskella*. Nematodes occur in both marine and terrestrial environments, and throughout their evolution have frequently transitioned between the two habitats [69]. Therefore, we initially suspected that marine nematodes were ancient hosts of *Lariskella,* potentially representing a link from the ancestrally marine hosts of Midichloriaceae to the primarily terrestrial hosts of *Lariskella*. However, the phylogenetic tree contradicts this notion, with the nematode-infecting *Lariskella* instead clustering among the insect-infecting *Lariskella* reference genomes (Fig 2). This placement, combined with the high nucleotide-level similarity with insect-infecting *Lariskella* strains, suggests the nematode population was likely infected by *Lariskella* relatively recently. Since the Thoracostomopsidae are primarily predators [70], and as predation can facilitate horizontal transmission of endosymbionts [71], we suspect the nematode population might have been infected through the ingestion of infected arthropods.

### *Lariskella* is not an obligate nutritional symbiont in nematodes

*Lariskella* has been hypothesized to play a nutritional role in ticks, acting as a potential source of B vitamins [29]. We confirmed the ability of *Lariskella* to synthesize several essential B vitamins, which could potentially support host nutrition. However, as *Lariskella* is absent in the majority of the nematode population, we can conclude that the nematode host does not rely on *Lariskella* to source essential B vitamins. The supplementation of B vitamins might instead be a method by which *Lariskella* could offset the fitness cost of infection, allowing it to spread in the host population more effectively, as has been shown in *Wolbachia* [83]. Though, notably *Lariskella* presence or absence has been shown to not affect the fitness of an insect host [31], suggesting *Lariskella* might not significantly contribute to nutrition.

### Microscopy confirms an intracellular localization

Microscopy confirmed the identity of the symbiont as *Lariskella*, with clear binding of the *Lariskella*-specific probe. The bacterial morphology in the TEM micrographs also resembled the *Lariskella* observed in insect cells [19], though the nematode-infecting *Lariskella* are notably more compact and less filamentous. Additionally, the microscopy demonstrated the intracellular localization of the *Lariskella* symbionts, confirming *Lariskella* represents a true intracellular infection of the nematodes and did not originate from external contamination. Furthermore, both the TEM and DOPE-FISH microscopy confirmed *Lariskella* colonised the developing oocytes within the gravid females, strongly suggesting vertical transmission of *Lariskella* from mother to offspring. Additionally, all oocytes within infected females displayed abundant *Lariskella* signal, suggesting a high rate of transmission. However, despite this suspected transmission efficiency, the overall *Lariskella* infection rate in nematodes was suprisingly low. Most notably, male nematodes seemed to be entirely free of infection during DOPE-FISH microscopy (Supplementary figure S6). While some *Lariskella* probe signal could be observed within males (Supplementary figure S7), these signals did not form clear cell-shaped structures or colocalized with the DAPI stain, and thus most likely originated from incomplete washing of unbound probe. This was confirmed with diagnostic PCR which also failed to detect *Lariskella* within male nematodes.

### The observed *Lariskella* infection rate might suggest reproductive manipulation

The overall infection rate of the nematode was found to be approximately 20%. This is far lower than infection rates typically reported for *Lariskella,* with studies reporting *Lariskella* frequencies around 47%-52% in ticks [42,72,73], 77%-100% in *Nysius* stinkbugs [19], 66%-100% in leaf-footed bugs [31], and 100% in psyllids [18]. Additionally, our results suggest that all male nematodes were free of *Lariskella* infection. A reduced prevalence of *Lariskella* in males is a known phenomenon. Female ticks have abundant *Lariskella* infections, while males only harbor small to negligible numbers of *Lariskella* [16,28,74]. Becker et al. (2023) suggested that *Lariskella* might be maternally transferred to all offspring, but that the *Lariskella* titer diminishes in males over the course of their development [28], similar to what has been observed for the related bacteria *Midichloria mitochondrii* [75]. However, while low in abundance, male ticks still harbor *Lariskella*, and *Lariskella* sequences have been detected in males by both qPCR [42] and amplicon sequencing [28,74]. In contrast, we failed to detect any *Lariskella* within males using diagnostic PCR or DOPE-FISH microscopy. Notably, the DOPE-FISH microscopy was sensitive enough to visualize individual *Lariskella* cells and still failed to detect any *Lariskella* infection in males.

Our observed lack of male infection led us to carefully hypothesize that *Lariskella* might be a reproductive parasite in nematodes. *Lariskella* is a reproductive manipulator in insects, where it has recently been demonstrated to be capable of inducing cytoplasmic incompatibility [31]. However, cytoplasmic incompatibility relies on infected males, and our results suggest an absence of infected males in the nematode population. Instead, the infection pattern observed in nematodes appears indicative of a male-killing or feminizing effect, as this would explain both the lack of male infection and the lower male prevalence. Additionally, a feminizing or male-killing effect would be more in line with the observed infection rate. As mentioned above, *Lariskella* tends to occur at higher frequencies in arthropods, which is considered typical of symbionts inducing cytoplasmic incompatibility [76]. In contrast, feminizing or male-killing symbionts are typically maintained at lower frequencies [6], to allow for sufficient male births to sustain the population. Our observed female infection rate (26%) appears in line with those observed for feminizing symbionts (15%-42%) [77–79] or male-killing symbionts (<40%) [6]. We also found potential orthologs of putative male-killing genes within the three analyzed *Lariskella* genomes (Supplementary file S1A). The inferred orthology was found to be based on a shared Cro/C1-type helix-turn-helix DNA-binding domain, which is commonly encountered in putative male-killing genes (Supplementary figure S8) [80–82]. However, attempts to culture the nematode have failed, and we thus lack the ability to experimentally test for male-killing or feminization.

## Conclusion

Here, we confirmed that *Lariskella* is an endosymbiont of a marine nematode species. Overall, this study illustrates that *Lariskella* is more widespread than previously described. With this novel discovery of *Lariskella* infecting nematodes, we posit *Lariskella* might also be a symbiont of terrestrial nematodes, as they would be more likely to be exposed to the known terrestrial hosts of *Lariskella.* Additionally, the infection of a marine host demonstrates that a marine habitat shift is no obstacle to *Lariskella,* and it could potentially infect other aquatic hosts. We suggest a more thorough sampling of marine invertebrates and terrestrial nematodes is needed, and suspect it would likely reveal a more widespread distribution of *Lariskella.*

## Acknowledgements

We would like to acknowledge the Darwin Tree of Life Project, as their public *Lariskella* genomes were highly valuable to this study. We would also like to thank Gayun Kim for her assistance in the molecular lab work, as well as Molly Hunter, Edwin Umanzor, and Yu Matsuura for their helpful insights on *Lariskella*. We are grateful for the help and support provided by the Imaging, Sequencing, and Scientific Computing and Data Analysis sections of the Core Facilities at Okinawa Institute of Science and Technology Graduate University.

## Data availability

The data used in this study is publicly available on NCBI as BioProject PRJNA1272136. All commands and scripts used for the analyses are available from https://github.com/ArnoHagenbeek/Thoraco_Lariskella_paper_code

---

**Supplementary table S1:** Genome statistics of the *Lariskella* MAG and reference genomes.

|  | *Lariskella* MAG from a Thoracostomopsidae nematode | Circularized *Lariskella* from *Epinotia ramella* (GCF_964019805.1) | Circularized *Lariskella* from *Hedychridium roseum* (GCF_964020185.1) |
| --- | --- | --- | --- |
| **Assembly size** | 1,506,628bp | 1,592,871bp | 1,585,685bp |
| **BUSCO score** | 92.9% complete<br>0% duplicated | 93.1% complete<br>0% duplicated | 92.6% complete<br>0.8% duplicated |
| **GC content** | 34.69% | 34.76% | 35.23% |
| **Contigs** | 321 | 1 | 1 |
| **CDS** | 1243 | 1448 | 1798 |
| **rRNA** | 1 | 3 | 3 |
| **tRNA** | 36 | 35 | 35 |

**Supplementary figure S1:**
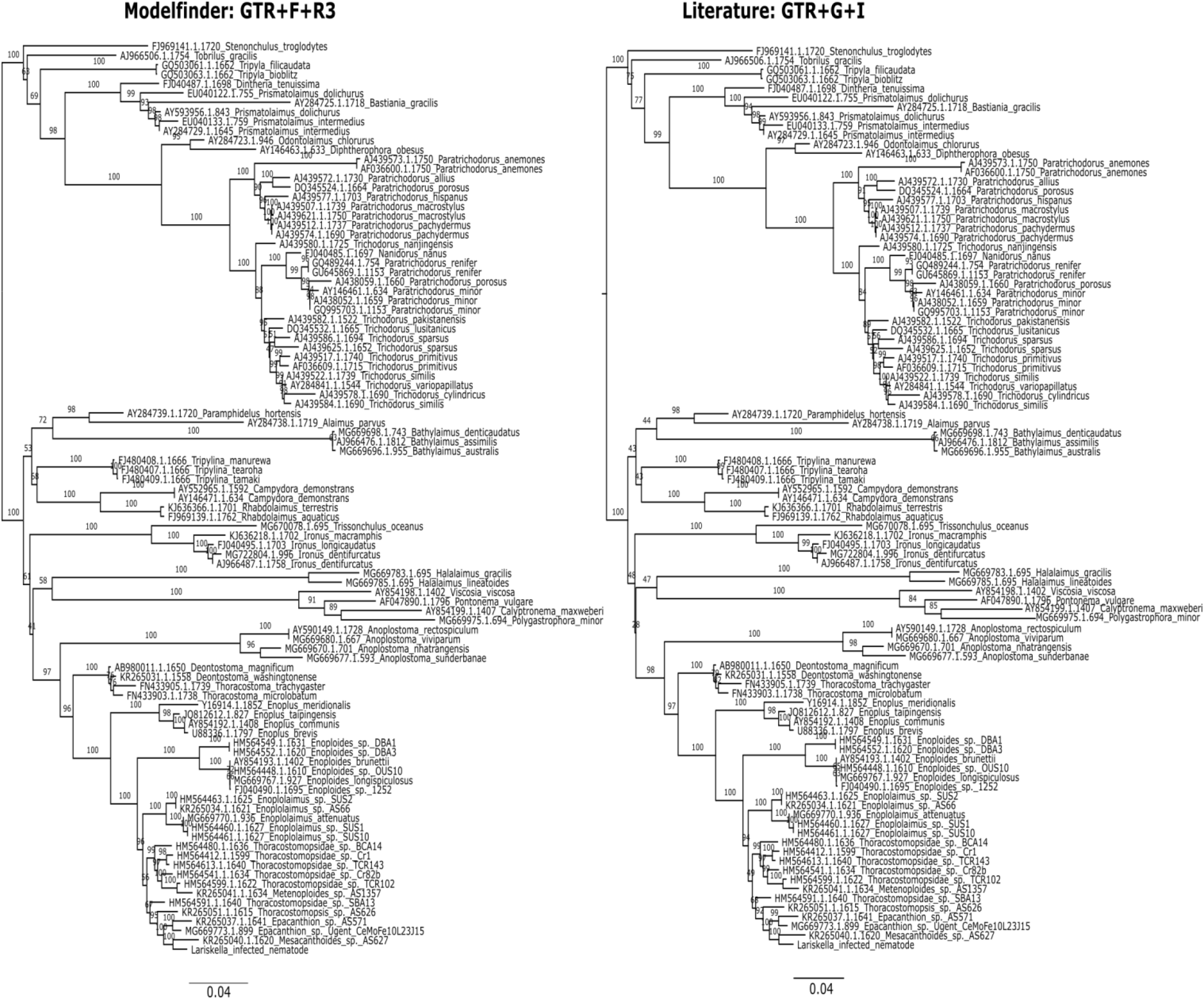
Comparison of different models for nematode phylogenetic tree reconstruction. Comparison of two phylogenetic trees reconstructed using the GTR+F+R3 model chosen by IQ-tree modelfinder, compared to the GTR+G+I model used in nematode phylogeny literature. The overall topology of the tree is the same, with only minor differences in branch lengths.

**Supplementary figure S2:**
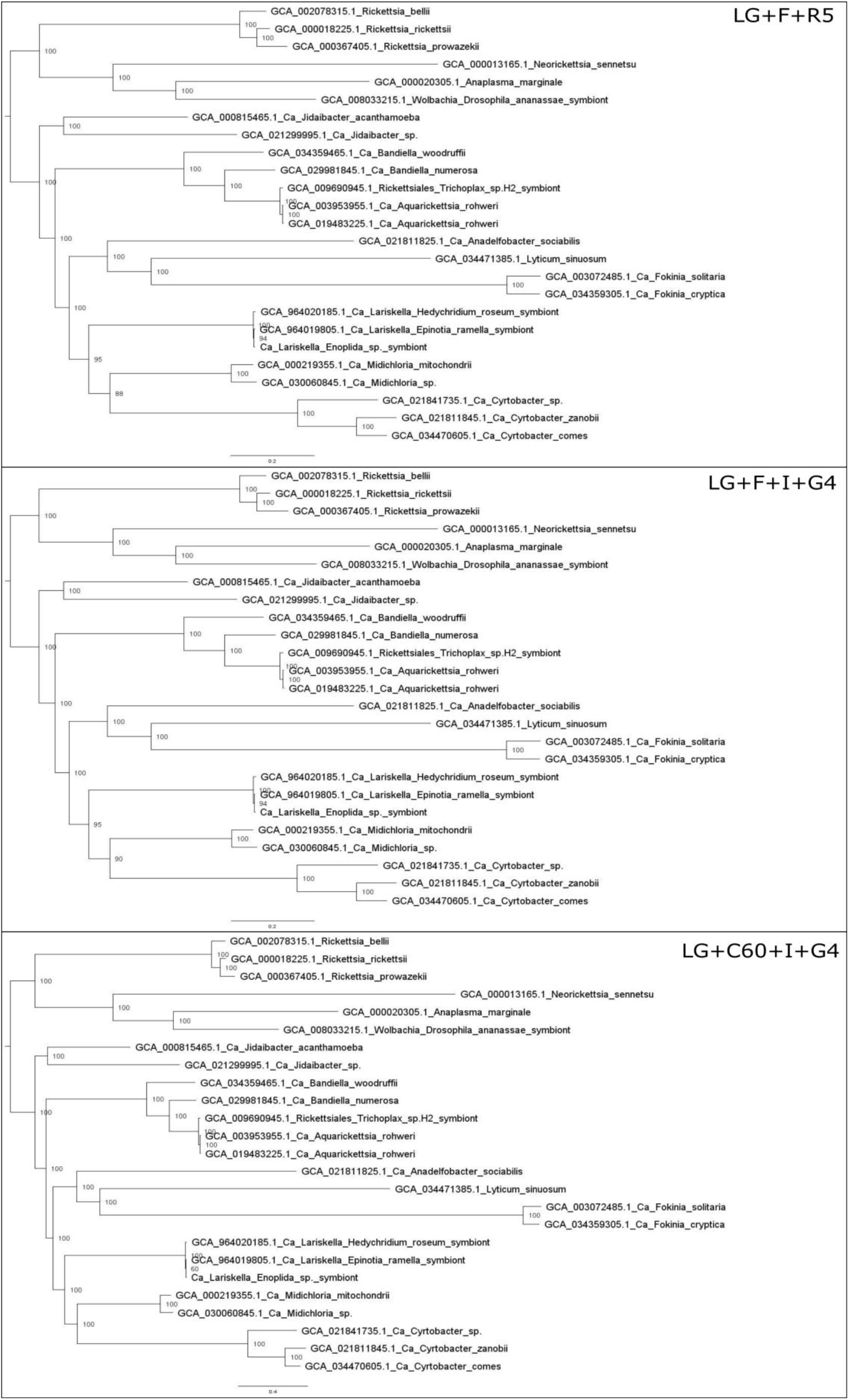
Comparison of different models for Midichloriaceae phylogenetic tree reconstruction. Comparison of different phylogenetic trees reconstructed using the LG+F+R5 model chosen by IQ-tree modelfinder, compared to the LG+F+I+G4 and LG+C60+I+G4 models. The overall topology of the tree is the same, with only minor differences in branch lengths.

**Supplementary figure S3:**
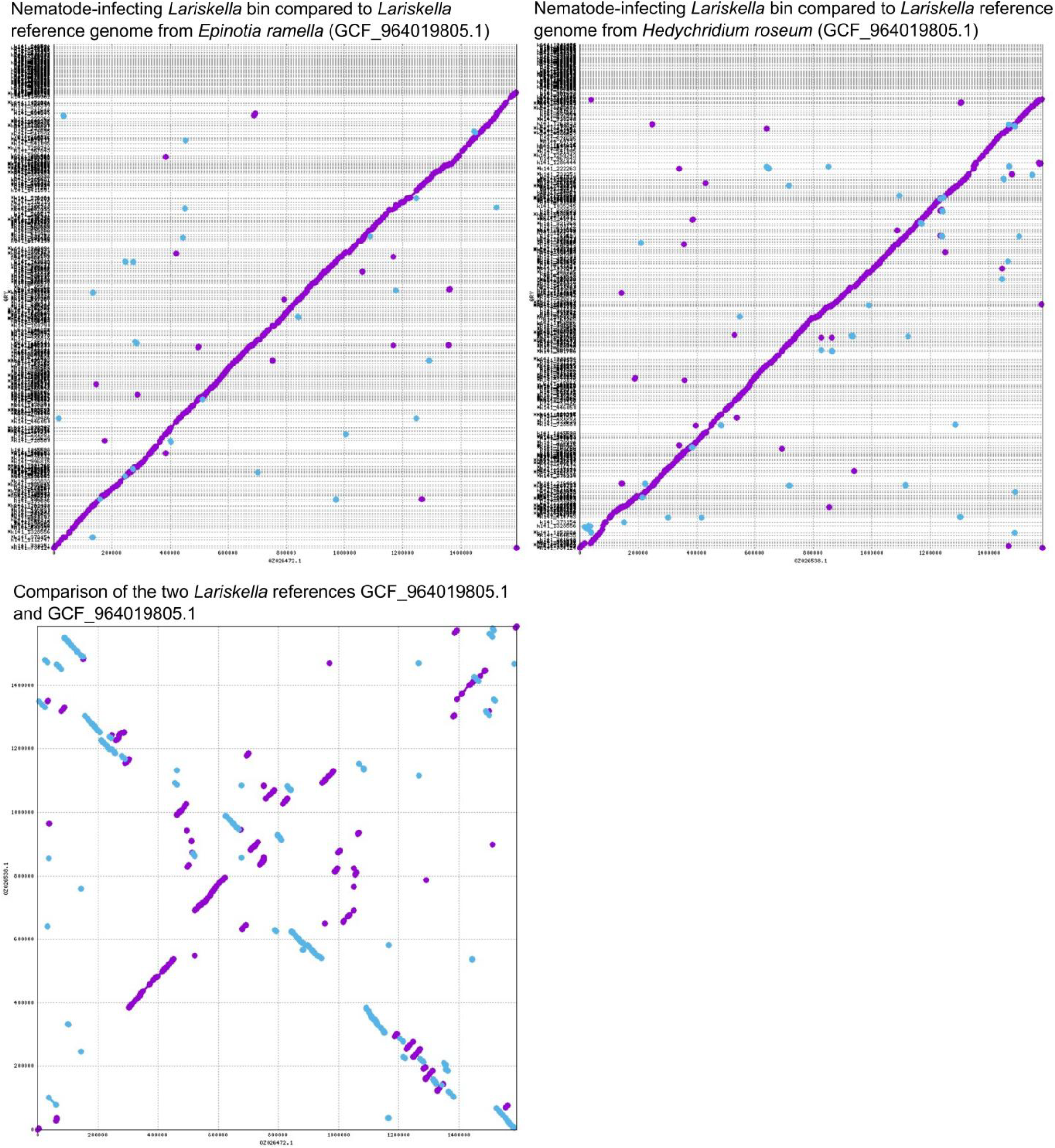
Mummer plots comparing the nucleotide sequences of the different *Lariskella* draft and reference genomes. Mummer plots based on Nucmer alignments, comparing the various *Lariskella* genomes to each other. Due to the fragmentation present in the nematode *Larikella* MAG, rearrangements are not visible. However, a high degree of nucleotide-level similarity is apparent.

**Supplementary figure S4:**
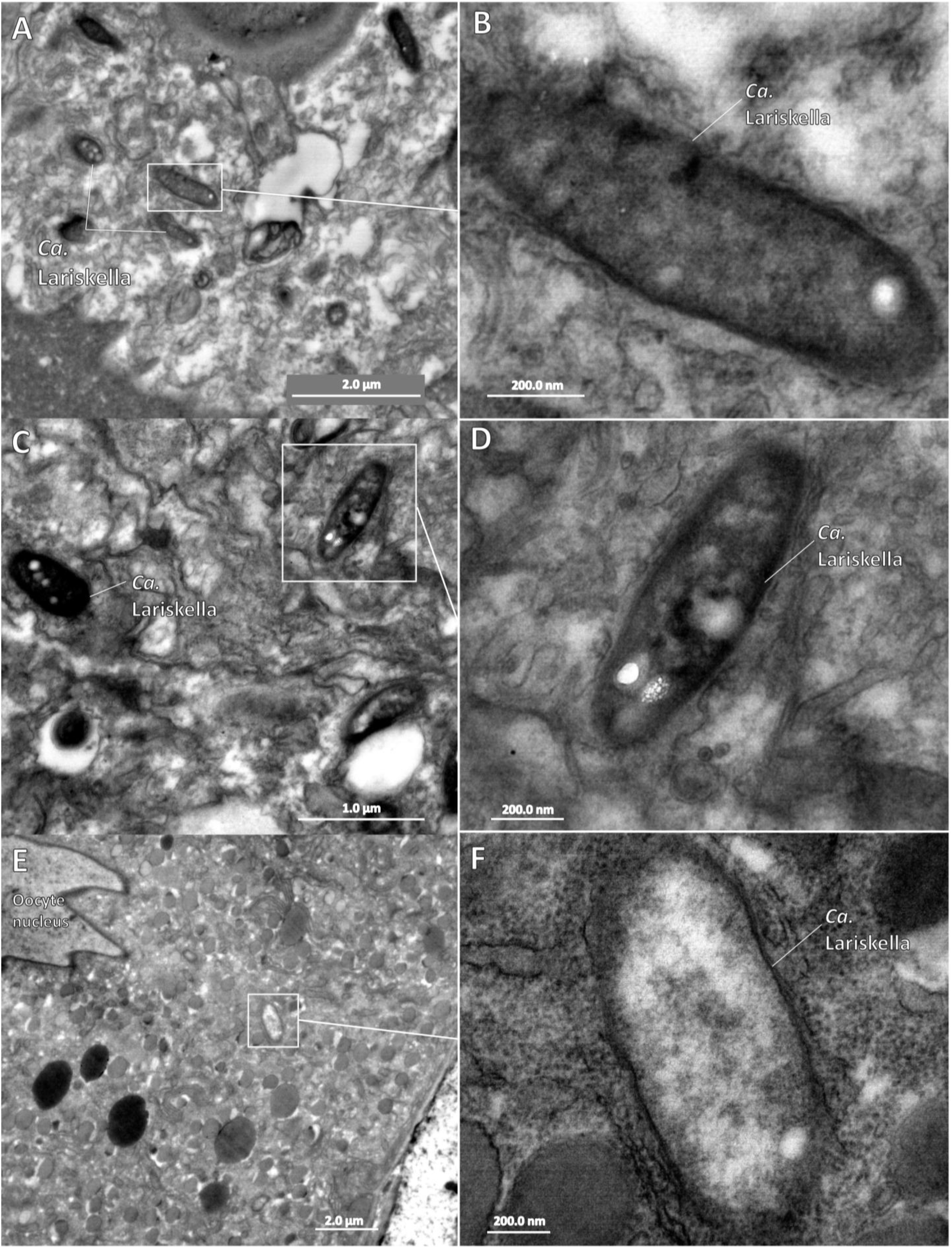
Supplementary TEM micrographs. Additional TEM micrographs displaying *Lariskella* cells within somatic tissue (A to D) and the oocyte (E and F)

**Supplementary figure S5:**
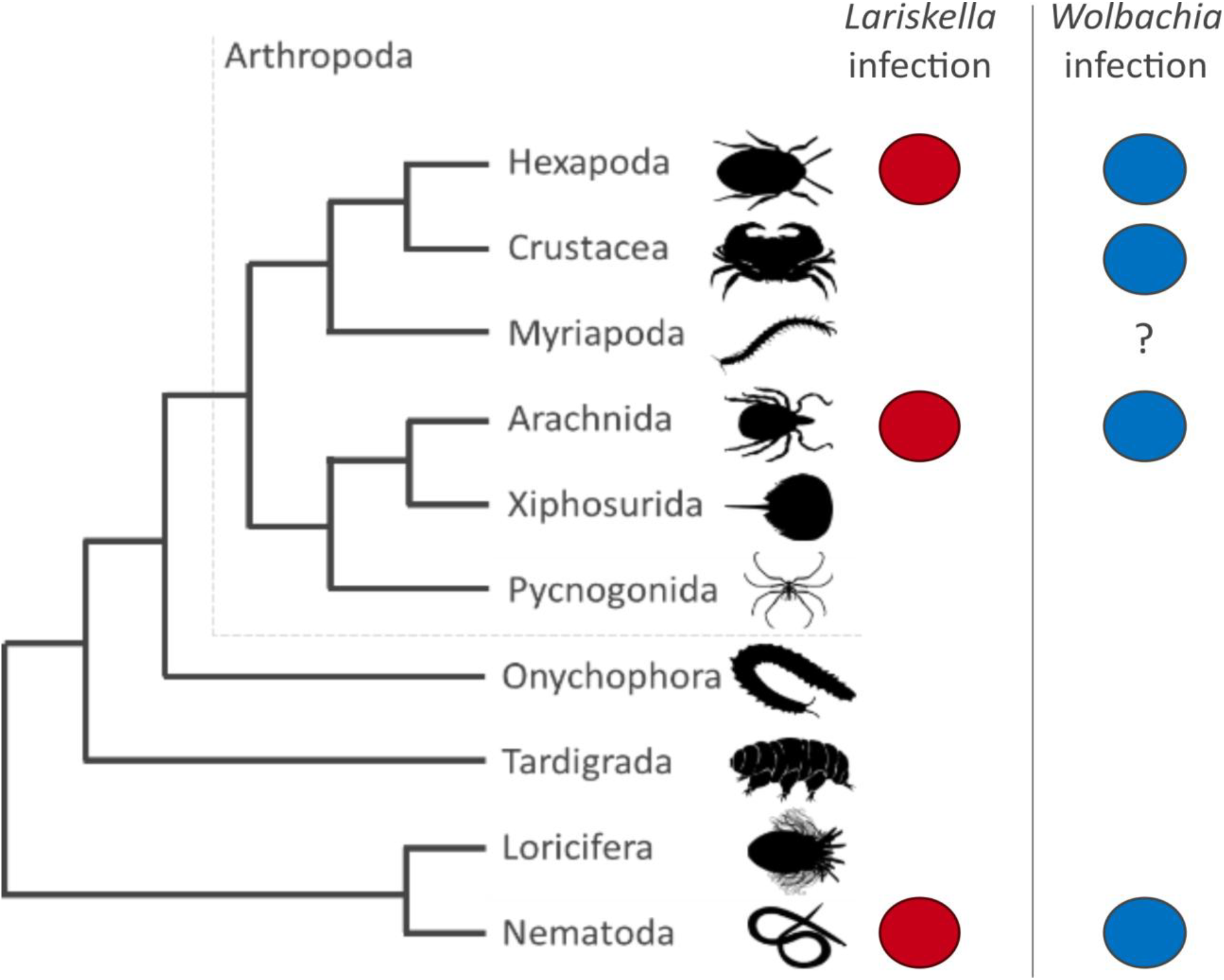
Dendrogram of Arthropoda, Nematoda, and related taxa. Dendrogram illustrating the similarities in the *Lariskella* and *Wolbachia* host ranges. Blue circles illustrate that Wolbachia is known to infect this taxon. A red circle illustrates that *Lariskella* has been reported to infect this taxon. Crustacea are the only known *Wolbachia* hosts for which no *Lariskella* has been found. There are also a few reports of *Wolbachia* in Myriapoda, but this presence is not well-documented, and we thus denoted the infection status with a question mark.

**Supplementary figure S6:**
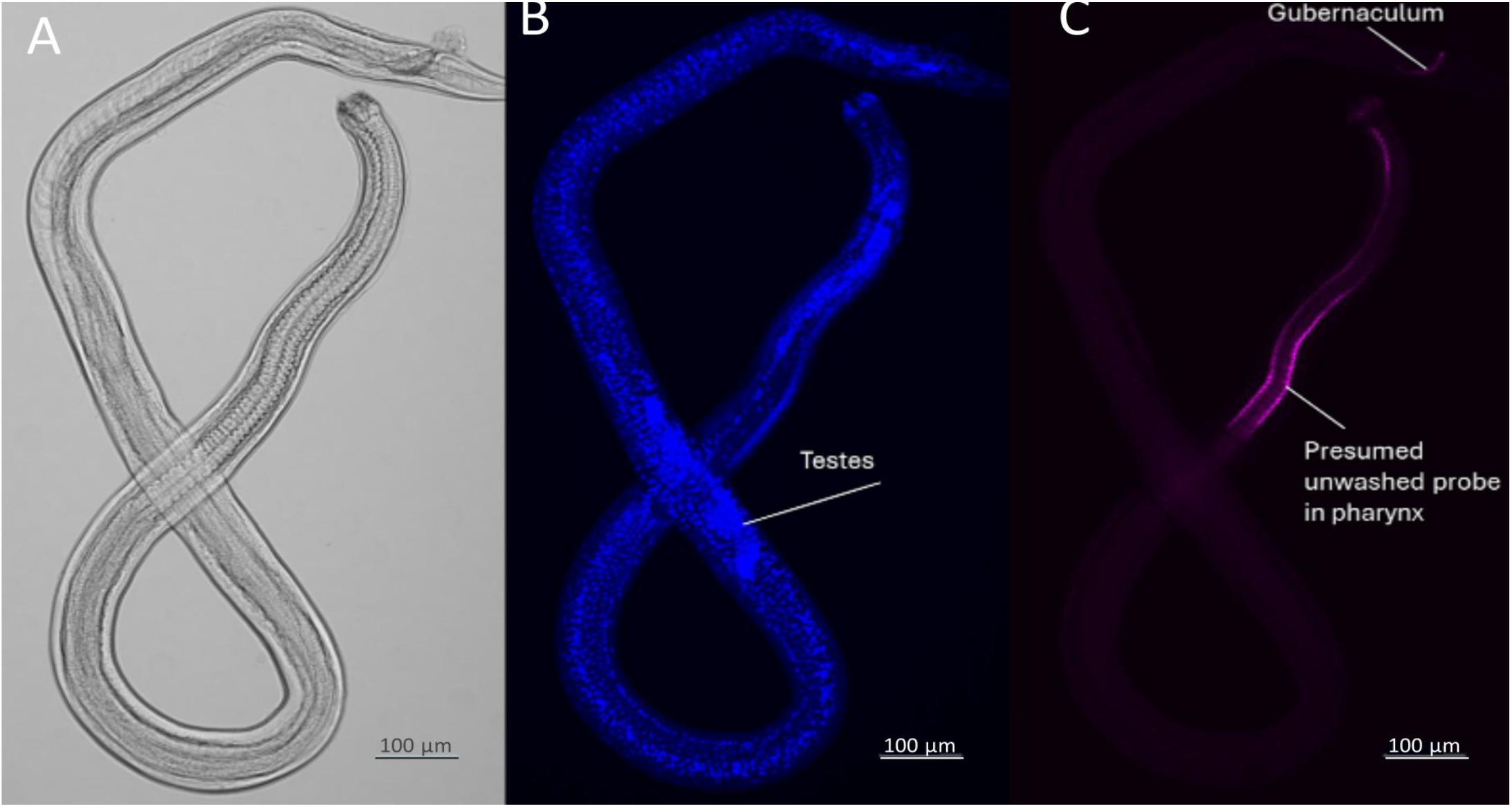
DOPE-FISH images of a whole male nematode. A) bright field image of the male nematode. B) DAPI channel of the male nematode DOPE-FISH, showing the distribution of nuclei throughout the nematode. C) *Lariskella*-probe channel of the DOPE-FISH, showing signal within the pharyngeal tissue of the male nematode. This signal did not form cells, and was commonly observed even within uninfected nematodes, suggesting it originates from probe that did not sufficiently wash out.

**Supplementary figure S7:**
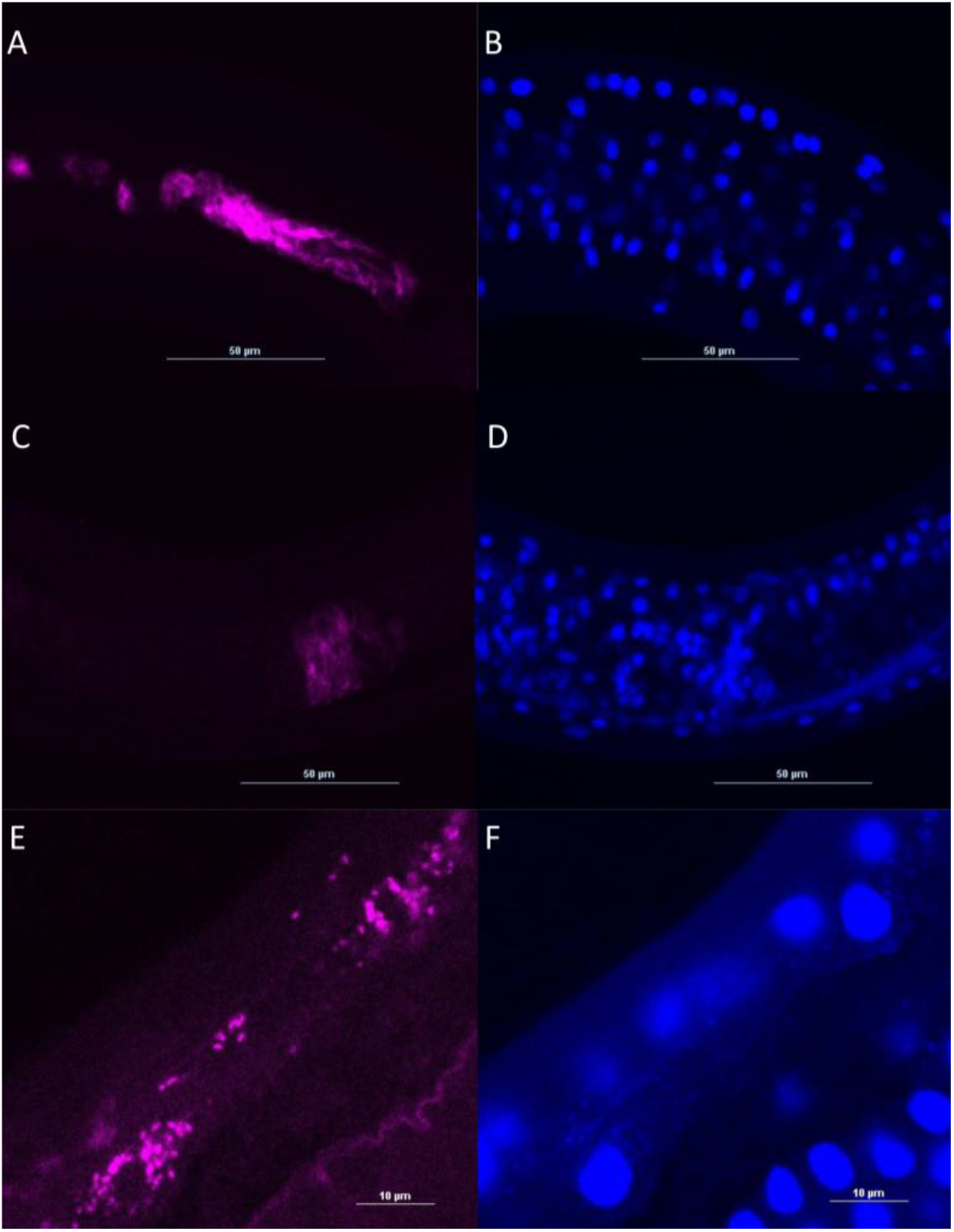
DOPE-FISH images of *Lariskella* probe signal in males. A to D) DOPE-FISH of male nematode *Lariskella* signal. A,C) *Lariskella* probe channel showing the probe does not locate in cell-like structures. B,E) DAPI channel showing DAPI does not co-locate with the *Lariskella* signal. E, F) DOPE-FISH of infected female nematode. E) *Lariskella* probe channel displaying that the probe signal forms into clear cell-like structures. F) DAPI channel showing weak DAPI signal co-locating with the cells indicated by the *Lariskella*-probe signal.

**Supplementary figure S8:**
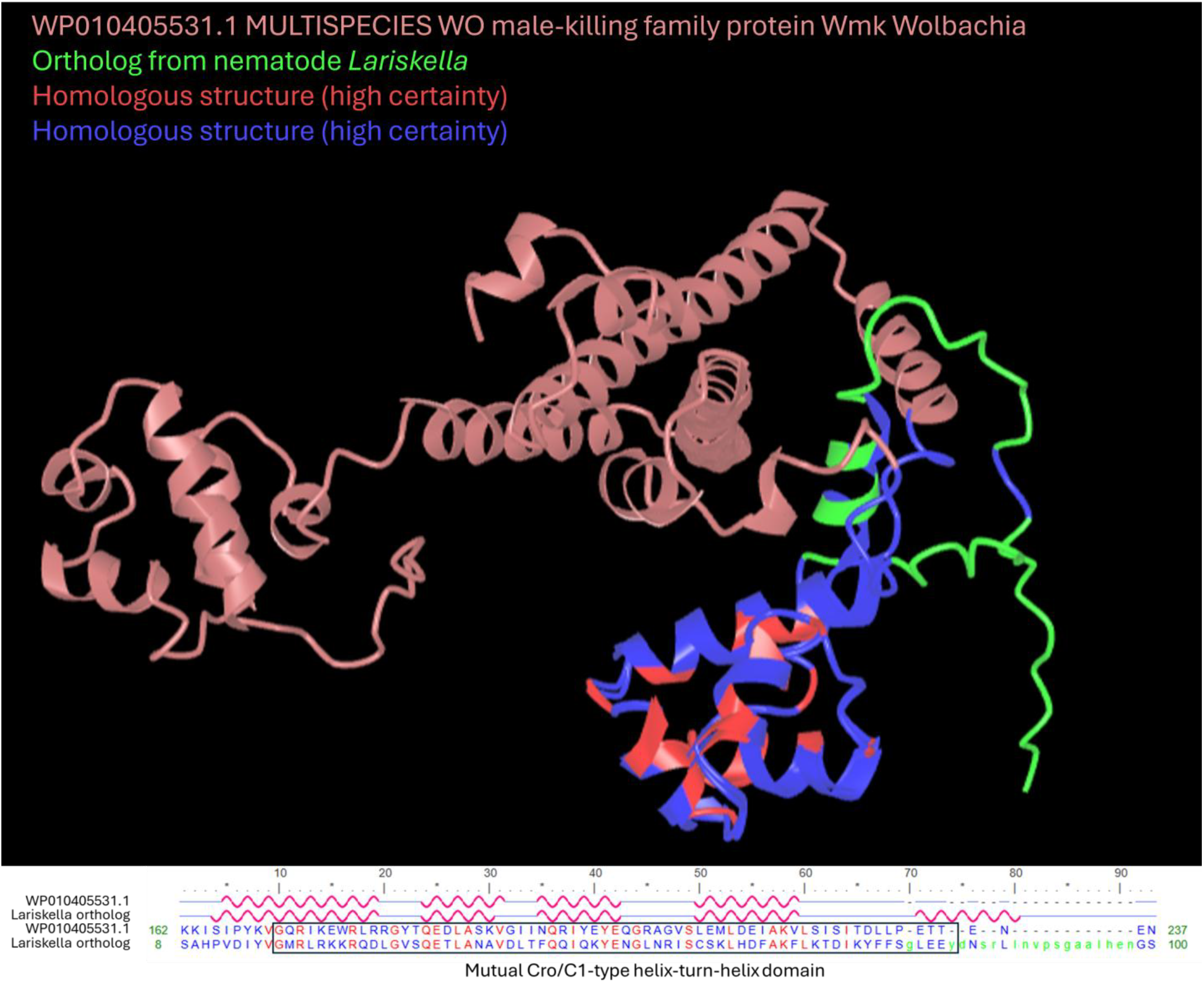
Structural alignment of putative *wmk* ortholog structures. Three-dimensional alignment of a putative *Wolbachia wmk* gene with a putative *Lariskella wmk* ortholog. Orthology was based on orthofinder and BLASTp searches. Blue and red coloration indicates homologous structures shared by both proteins, and correspond to the blue and red letters in the protein sequence below the image. The bracketed protein sequence indicates the CRO/C1 domain found in both proteins by Interproscan.

**Supplementary file S1A: Protein sequence fasta of the putative *Lariskella wmk* orthologs**

>GCA_964019805.1_Lariskella_in_Epinotia.fna_00359 hypothetical protein

MSKKKATSAHPVDIYVGMRLRKKRQDLGVSQETLANAVDLTFQQIQKYEKGLNRISCSKL HDVAKFLKTDIKYFFSGLEEYDNSRLLNVPSGAALHENGSQYKSHNSDFAELQDAFDSIK

QKELKKSIINIVQNLALN

>GCA_964020185.1_Lariskella_in_Hedychridium.fna_00536 hypothetical protein MSKKKATSAHPVDIYVGMRLRKKRQDLGVSQETLANAVDLTFQQIQKYEKGLNRISCSKL

HDFAKFLKTDIKYFFSGLEEYDNSRLLNVPSGAALH

>Lariskella_in_nematode.fna_00135 hypothetical protein

MSKKKATSAHPVDIYVGMRLRKKRQDLGVSQETLANAVDLTFQQIQKYENGLNRISCSKL

HDFAKFLKTDIKYFFSGLEEYDNSRLLNVPSGAALHENGSQYKSHNSDFAE

**Supplementary file S1B: Protein sequence fasta of the *wmk* reference sequences used to infer orthology**

>MK873003.1_1 Wolbachia endosymbiont of Drosophila borealis putative DNA-binding protein (WD0626_wmk-like) gene, complete cds VRKENNCSSFLDYKAIGQEVRNRRLAKRYTQKDLAGKIGVTYQIVLQYEKGTRKISIEKL

YAIAEALSVSIVDLIPVSNEKIYLENKEEEILNLIREYKKISGQELRKVFCLLTKFARVG

EENSKKTERIKIAKGLVKEGISIDIVSKTIGLSADECVEEKVGSIYCKIGKKIKEWRIVR

GYTQKDLAEKMSTTRDEISNYEQGRVAIPLEKLYAIAETLSINIMDLLELTEDADDKVEN

ELPNLIEEYKEIESQELRYVLIKSLLEGIQIYEEKVKRAEKMKIAKDLVKKGISTDIILQ

ITGLSLDEIQQV

>MK873082.1_1 Wolbachia endosymbiont of Drosophila innubila isolate wInn_0626 hypothetical protein gene, partial cds VRKENNCSSFLDYKAIGQEVRNRRLAKRYTQKDLAGKIGVTYQIVLQYEKGTRKISIEKL

YAIAEALSVSIVDLIPVSNEKIYLENKEEEILNLIREYKKISGQELRKVFCLLTKFARVG

EENSKKTERIKIAKGLVKEGISIDIVSKTIGLSADECVEEKVGSIYCKIGKKIKEWRIVR

GYTQKDLAEKMSTTRDEISNYEQGRVAIPLEKLYAIAETLSINIMDLLELTEDADDKVEN

ELPNLIEEYKEIESQELRYVLIKSLLEGIQIYEEKVKRAEKMKIAKDLVKKGISTDIILQ

ITGLSLDEIQQV

>AAS14326.1 transcriptional regulator, putative [Wolbachia endosymbiont of Drosophila melanogaster]

MANISIRYKIAQKVRSWRLKRGYTQKDLAGKIGVTYQVVLQYEKGTRKISIEKLYAIAEVLSVGIIDLIP

VSSEKICLKNEEEEILNLVRKYKTINDQELRKVFYLLTKFTRVGEKSSKKAEKVKIAKGMVKAGISVDIV

SQAIGLSANECVEEKTGSIYYQIGKKIKEWRLVREYTQKDLAEKMDTTRDEISNYEQGRVAIPLEKLYAI

AETLSISITDLLIEEDEIVESELPDLIKEYKKIESQELRYALIKSLFESIQICEEKVKRAEKMKIAKDLV

KGGISTDIILQITGLSLGEIQQI

>AAS14324.1 transcriptional regulator, putative [Wolbachia endosymbiont of Drosophila melanogaster]

MVLFVEKSLDYKVGEKLKSWRLERGYTQKDLAEKLGVKYWVILQYEKGNRRISIERLYAITEALSISITD

LIPISKSCLEDEGEEILNLVREYKKINDQELRKMFCLLTNFVQVSEKSSKKAEKIKIAKGLVKAGVSVDI

VAKTIGLSADECVEEKGGSIYCQIGKKIKEWRLVREYTQKDLVEKMSTTRDEISNYEQGRTAVPLDKLYE

MAEALSINITDLLIKEGSKVKNELPDLIKEYKEIESQELRHALIKSLFEGIRICEEKVREIERIKVAKDL VKGGISIDIILQIIGLSVDQIA

>MK873081.1_1 Wolbachia endosymbiont of Drosophila innubila isolate wInn_0623 hypothetical protein gene, partial cds

KINDQELRRMFCLLTKFVKVSEKSSKKSEKIKIAKGLVKAGVSVDIVAKTIGLPADECIE

EKVGSIYCQIGKKIKEWRLVREYTQKDLAEKMNTTRDEISNYEQGRVATPLGKLYEIAEA

LSISITDLLTEEDEGSRVENELPDLIKEYKEIESQELRNALIKSLFEGIRICEEKVREIE

RIKVAKDLVKGGISIDIILQAVGLPVDIVLDG

>MK873002.1_1 Wolbachia endosymbiont of Drosophila borealis putative DNA-binding protein (WD0623-like) gene, complete cds

VEKILDYEVGEKVKSWRLERGYTQKDLAEKIGVKYWVILQYEKGNRRISIKRLYAIAGAL

SVSITDLITASKEKIGFKNEEGEILNLVREYKKINDQELRRMFCLLTKFVKVSEKSSKKS

EKIKIAKGLVKAGVSVDIVAKTIGLPADECIEEKVGSIYCQIGKKIKEWRLVREYTQKDL

AEKMNTTRDEISNYEQGRVATPLGKLYEIAEALSISITDLLTEEDEGSRVENELPDLIKE

YKEIESQELRNALIKSLFEGIRICEEKVREIERIKVAKDLVKGGISIDIILQAVGLPVDI VLDG

>AAS13997.1 transcriptional regulator, putative [Wolbachia endosymbiont of Drosophila melanogaster]

MVVFVEKSLDCKVGEKVKNWRLERGYTQKDLAEKIGVKYWVILQYEKGNRGISIKRLYAIAEALSVSITN

LIPASKEKIGFKNEEGEILNLVREYKKINDHELRRMFCLLTKFVQVSEKSSRKSEKIKIANGLVKAGISV

DIVSKTIGLSADECIEEKVGSIYYKIGKKIKEWRLVREYTQKDLGEKMSTTRHEVSNYEQGRTAVPLDKL

YEMAEALSINITDLLIEKDEGSRVENELPNLIKEYKEIESQELRNALIKSLFEGIRICEEKVREIERIKV AKDLVKGGISIDIILQAVGLPVDIVLDR

>MK873005.1_1 Wolbachia endosymbiont of Drosophila bifasciata putative DNA-binding protein (WD0626_wmk-like) gene, complete cds

MFVLQLSENMSIEKKIDPDSLCYYITQKVKNVRFELNCTQGEFAEKTGVAKSLIGKYERG

IHSILPKTLEDMAKKLSKDIENFFPESTNCISSEDKKLFDLVQTLKRIKDWKVRDTVCVL

TRFLSEGIQISKVTEEIRPISYQMVQKAKDWRFARGRTQMEVADKSGMPHGQVVRYEKGE

VSFKIAPKMAKGLSLHYGVLLPTSKAERYCEDEDGEGEKKILSIMREYQKIDNQKLKDLW

YSFLSEVVKISEEKDS

>MK873080.1_1 Wolbachia endosymbiont of Drosophila innubila isolate wInn_0622 hypothetical protein gene, partial cds

VVREYKDEDEEIFYLTKIYENQKLGKIVPSLVRFVHISEKINQEEARLEVAKNLVKEGVS

VDIISQATGLSIYEYDNKEREICTDSIYYRIXXXXREWRLIRRYTQKDLADKVGLTLKEI

HEYEIGYTAISFDKLYQIAEGLSVNIKVLLPKTNEDSELLNLLRKTEEQELVKKFLSRDM

KNSKEKVKKTEKIKVAKNLVKAGVSTDVILRASGLTADECEN

>MK873001.1_1 Wolbachia endosymbiont of Drosophila borealis putative DNA-binding protein (WD0622-like) gene, complete cds

MFVSASNVSYGIGQKIENCRLMRGHTQIGLASQIGLTYKEVNSYENGYISIPIEVLYTIA

KELSVSVTDLLPESVVVREYKDEDEEIFYLTKIYENQKLGKIVPSLVRFVHISEKINQEE

ARLEVAKNLVKEGVSVDIISQATGLSIYEYDNKEREICTDSIYYRIGQRIREWRLIRRYT

QKDLADKVGLTLKEIHEYEIGYTAISFDKLYQIAEGLSVNIKVLLPKTNEDSELLNLLRK

TEEQELVKKFLSRDMKNSKEKVKKTEKIKVAKNLVKAGVSTDVILRASGLTADECEN

>AAS14323.1 transcriptional regulator, putative [Wolbachia endosymbiont of Drosophila melanogaster]

MFVSVSDISSISYKIGQKIEDSRLMRGHTQVELASEIGLTYQEVNSYENGYIPIPIEVLYVIARVLSVNA

IDLLPEPVIVREDSYEDEEILYLTKIYENQKLGKIVPSLVRFVHISEKINQEEARLEIAKNLVKEGVSVD

IISQATGLSIYEYDNTEREICTDSIYYRIGQRIREWRLIRRYTQKDLADKVGVTLKEIHEYERGYTTILF

DKLYEIAGALSVNIKVLLPETRESKKLLSLINEYREPESLDALVKSLSEDMKSGKEKVKKAEKIRVAKNL

AKAGVAIDIIVRASGLTADECEN

>WP_015587820.1 MULTISPECIES: WO male-killing family protein Wmk [unclassified Wolbachia]

MKKENKCSNFLDYKVIGQEVRNRRLAKGYTQKDLAKKIGTTYQVILQYEKGTRRISIKKLYELAEALSTT

VRDLACGQEVSNEKGYEGEEVLNLVRRHKEIKNQELRETFYLLTKFIRIGEEESGKVVKIEVAKGLVKEG

VSAHVISQTTSLSIDEYDNDEKKISIPYKVGQRIKEWRLRRGYTQEDLASKVGIINQRIYEYEQGRAAVS

LEMLDEIAKVLLINITDLLPETRENENSEAELSKLIEEYKKIKSQELRHVLIKSLFESIQVCKEKVKRVE

KMKIAKNLVKEGISINIILKTVGISLDEIQQI

>WP_010405531.1 MULTISPECIES: WO male-killing family protein Wmk [Wolbachia]

MKKENKCSNFLDYKVIGQEVRNRRLAKGYTQKDLAKKIDTTYQVILQYEKGTRRISIKKLYELAEALSTT

ARDLACGQEVSNEERYEEEEILNLVRRHKEIKDQELRETFYLLTKFIRISEEESGKAVKVEVAKGLVKEG

VSAHVISQTTSLSIDEYDNDEKKISIPYKVGQRIKEWRLRRGYTQEDLASKVGIINQRIYEYEQGRAGVS

LEMLDEIAKVLSISITDLLPETTENENSEVELSRLIEEYKKIKSQELRHVLIKSLFESIQVCKEKVKRVE

KMKIAKNLVKEGISINIILKTVGISLDEIQQI

>AAS14223.1 transcriptional regulator, putative [Wolbachia endosymbiont of Drosophila melanogaster]

MKKENKCSNFLDYKVIGQEVRNRRLAKGYTQKDLAKKIDTTYQVILQYEKGTRRISIKKLYELAEALSTT

ARDLACGQEVSNEERYEEEEILNLVRRHKEIKDQELRETFYLLTKFIRISEEESGKAVKVEVAKGLVKEG

VSAHVISQTTSLSIDEYDNDEKKISIPYKVGQRIKEWRLRRGYTQEDLASKVGIINQRIYEYEQGRAAVS

LEMLNEIAKVLLINITDLLPETRENENSEAELSRLIEEYKKIKSQELRDVLIKSLLESIQICKEKVKKIE

KIKIAKNLVKEGISINIILKTVGISLDEIQQI

## References

1. Ettema TJG, Andersson SGE. The α-proteobacteria: the Darwin finches of the bacterial world. Biol Lett 2009;5:429–32.

2. Renvoisé A, Merhej V, Georgiades K et al. Intracellular Rickettsiales: Insights into manipulators of eukaryotic cells. Trends Mol Med 2011;17:573–83.

3. Hosokawa T, Koga R, Kikuchi Y et al. Wolbachia as a bacteriocyte-associated nutritional mutualist. Proc Natl Acad Sci 2010;107:769–74.

4. Zhao D, Zhang Z, Niu H et al. Pathogens are an important driving force for the rapid spread of symbionts in an insect host. Nat Ecol Evol 2023;7:1667–81.

5. Pannebakker BA, Loppin B, Elemans CPH et al. Parasitic inhibition of cell death facilitates symbiosis. Proc Natl Acad Sci U S A 2007;104:213–5.

6. Engelstädter J, Hurst GDD. The Ecology and evolution of microbes that manipulate host reproduction. Annu Rev Ecol Evol Syst 2009;40:127–49.

7. Montagna M, Sassera D, Epis S et al. “Candidatus Midichloriaceae” fam. nov. (Rickettsiales), an ecologically widespread clade of intracellular Alphaproteobacteria. Appl Environ Microbiol 2013;79:3241–8.

8. Giannotti D, Boscaro V, Husnik F et al. The “other” Rickettsiales: an overview of the family “Candidatus Midichloriaceae.” Appl Environ Microbiol 2022;88:e02432–21.

9. Vannini C, Ferrantini F, Schleifer K-H et al. “Candidatus Anadelfobacter veles” and “Candidatus Cyrtobacter comes,” two new Rickettsiales species hosted by the protist ciliate Euplotes harpa (Ciliophora, Spirotrichea). Appl Environ Microbiol 2010;76:4047–54.

10. Szokoli F, Sabaneyeva E, Castelli M et al. “Candidatus Fokinia solitaria”, a novel “stand-alone” symbiotic lineage of Midichloriaceae (Rickettsiales). PLOS ONE 2016;11:e0145743.

11. Schulz F, Martijn J, Wascher F et al. A Rickettsiales symbiont of amoebae with ancient features. Environ Microbiol 2016;18:2326–42.

12. Muñoz-Gómez SA, Hess S, Burger G et al. An updated phylogeny of the Alphaproteobacteria reveals that the parasitic Rickettsiales and Holosporales have independent origins. Rokas A, Wittkopp PJ, Irisarri I (eds.). eLife 2019;8:e42535.

13. Driscoll T, Gillespie JJ, Nordberg EK et al. Bacterial DNA sifted from the Trichoplax adhaerens (Animalia: Placozoa) genome project reveals a putative Rickettsial endosymbiont. Genome Biol Evol 2013;5:621–45.

14. Kamm K, Osigus H-J, Stadler PF et al. Genome analyses of a placozoan rickettsial endosymbiont show a combination of mutualistic and parasitic traits. Sci Rep 2019;9:17561.

15. Klinges JG, Rosales SM, McMinds R et al. Phylogenetic, genomic, and biogeographic characterization of a novel and ubiquitous marine invertebrate-associated Rickettsiales parasite, Candidatus Aquarickettsia rohweri, gen. nov., sp. nov. ISME J 2019;13:2938–53.

16. Kurilshikov A, Livanova NN, Fomenko NV et al. Comparative metagenomic profiling of symbiotic bacterial Communities associated with Ixodes persulcatus, Ixodes pavlovskyi and Dermacentor reticulatus ticks. PLOS ONE 2015;10:e0131413.

17. Toju H, Tanabe AS, Notsu Y et al. Diversification of endosymbiosis: replacements, co-speciation and promiscuity of bacteriocyte symbionts in weevils. ISME J 2013;7:1378–90.

18. Morrow JL, Hall AAG, Riegler M. Symbionts in waiting: the dynamics of incipient endosymbiont complementation and replacement in minimal bacterial communities of psyllids. Microbiome 2017;5:58.

19. Matsuura Y, Kikuchi Y, Meng XY et al. Novel clade of Alphaproteobacterial endosymbionts associated with stinkbugs and other arthropods. Appl Environ Microbiol 2012;78:4149–56.

20. Mancini E, Sabatelli S, Hu Y et al. Uncovering active bacterial symbionts in three species of pollen-feeding beetles (Nitidulidae: Meligethinae). Microb Ecol 2023;85:335–9.

21. Sassera D, Beninati T, Bandi C et al. ‘Candidatus Midichloria mitochondrii’, an endosymbiont of the tick Ixodes ricinus with a unique intramitochondrial lifestyle. Int J Syst Evol Microbiol 2006;56:2535–40.

22. Mediannikov OI, Ivanov LI, Nishikawa M et al. Microorganism “Montezuma” of the order Rickettsiales: the potential causative agent of tick-borne disease in the far east of Russia. Zh Mikrobiol Epidemiol Immunobiol 2004:7–13.

23. Bazzocchi C, Mariconti M, Sassera D et al. Molecular and serological evidence for the circulation of the tick symbiont Midichloria (Rickettsiales: Midichloriaceae) in different mammalian species. Parasit Vectors 2013;6:350.

24. Mariconti M, Epis S, Gaibani P et al. Humans parasitized by the hard tick Ixodes ricinus are seropositive to Midichloria mitochondrii: is Midichloria a novel pathogen, or just a marker of tick bite? Pathog Glob Health 2012;106:391–6.

25. Mariconti M, Epis S, Sacchi L et al. A study on the presence of flagella in the order Rickettsiales: the case of ‘Candidatus Midichloria mitochondrii.’ Microbiology 2012;158:1677–83.

26. Sassera D, Lo N, Epis S et al. Phylogenomic evidence for the presence of a flagellum and cbb3 Oxidase in the free-living mitochondrial ancestor. Mol Biol Evol 2011;28:3285–96.

27. Grandi G, Chiappa G, Ullman K et al. Characterization of the bacterial microbiome of Swedish ticks through 16S rRNA amplicon sequencing of whole ticks and of individual tick organs. Parasit Vectors 2023;16:39.

28. Becker NS, Rollins RE, Stephens R et al. Candidatus Lariskella arthopodarum endosymbiont is the main factor differentiating the microbiome communities of female and male Borrelia-positive Ixodes persulcatus ticks. Ticks Tick-Borne Dis 2023;14:102183.

29. Buysse M, Duron O. Evidence that microbes identified as tick-borne pathogens are nutritional endosymbionts. Cell 2021;184:2259–60.

30. Lu M, Meng C, Zhang B et al. Prevalence of spotted fever group Rickettsia and Candidatus Lariskella in Multiple Tick Species from Guizhou Province, China. Biomolecules 2022;12:1701.

31. Umanzor EF, Kelly SE, Ravenscraft A et al. The facultative intracellular symbiont Lariskella is neutral for lifetime fitness and spreads through cytoplasmic incompatibility in the leaffooted bug, Leptoglossus zonatus. Front. Microbiol. 16:1595917. doi: 10.3389/fmicb.2025.1595917

32. Aivelo T, Tschirren B. Bacterial microbiota composition of a common ectoparasite of cavity-breeding birds, the hen flea Ceratophyllus gallinae. Ibis 2020;162:1088–92.

33. Gruber-Vodicka HR, Seah BKB, Pruesse E. phyloFlash: Rapid small-subunit rRNA profiling and targeted assembly from metagenomes. mSystems 2020;5:e00920–20.

34. Quast C, Pruesse E, Yilmaz P et al. The SILVA ribosomal RNA gene database project: improved data processing and web-based tools. Nucleic Acids Res 2013;41:D590–6.

35. Rozewicki J, Li S, Amada KM et al. MAFFT-DASH: integrated protein sequence and structural alignment. Nucleic Acids Res 2019;47:W5–10.

36. Minh BQ, Schmidt HA, Chernomor O et al. IQ-TREE 2: New models and efficient methods for phylogenetic inference in the genomic era. Mol Biol Evol 2020;37:1530–4.

37. Megen, Elsen, den van et al. A phylogenetic tree of nematode based on about 1200 full-length small subunit ribosomal DNA sequences. Nematol 11 2009 6 2009;11, DOI: 10.1163/156854109×456862.

38. Meldal BHM, Debenham NJ, De Ley P et al. An improved molecular phylogeny of the Nematoda with special emphasis on marine taxa. Mol Phylogenet Evol 2007;42:622–36.

39. Camacho C, Coulouris G, Avagyan V et al. BLAST+: architecture and applications. BMC Bioinformatics 2009;10:421.

40. Keeling PJ. Molecular phylogenetic position of Trichomitopsis termopsidis (Parabasalia) and evidence for the Trichomitopsiinae. Eur J Protistol 2002;38:279–86.

41. Deane J, Hill D, Brett S et al. Hanusia phi gen. et sp. nov. (Cryptophyceae): characterization of ‘Cryptomonas sp. Φ.’ Eur J Phycol 1998;33:149–54.

42. Mukhacheva TA, Kovalev SY. Bacteria of the family ‘Candidatus Midichloriaceae’ in sympatric zones of Ixodes ticks: Genetic evidence for vertical transmission. Microb Ecol 2017;74:185–93.

43. Madeira F, Madhusoodanan N, Lee J et al. The EMBL-EBI job dispatcher sequence analysis tools framework in 2024. Nucleic Acids Res 2024;52:W521–5.

44. Chen S. Ultrafast one-pass FASTQ data preprocessing, quality control, and deduplication using fastp. iMeta 2023;2:e107.

45. Li D, Liu C-M, Luo R et al. MEGAHIT: an ultra-fast single-node solution for large and complex metagenomics assembly via succinct de Bruijn graph. Bioinformatics 2015;31:1674–6.

46. Langmead B, Salzberg SL. Fast gapped-read alignment with Bowtie 2. Nat Methods 2012;9:357–9.

47. Danecek P, Bonfield JK, Liddle J et al. Twelve years of SAMtools and BCFtools. GigaScience 2021;10:giab008.

48. Kang DD, Li F, Kirton E et al. MetaBAT 2: an adaptive binning algorithm for robust and efficient genome reconstruction from metagenome assemblies. PeerJ 2019;7:e7359.

49. Alneberg J, Bjarnason BS, de Bruijn I et al. Binning metagenomic contigs by coverage and composition. Nat Methods 2014;11:1144–6.

50. Pan S, Zhao X-M, Coelho LP. SemiBin2: self-supervised contrastive learning leads to better MAGs for short- and long-read sequencing. Bioinformatics 2023;39:i21–9.

51. Sieber CMK, Probst AJ, Sharrar A et al. Recovery of genomes from metagenomes via a dereplication, aggregation and scoring strategy. Nat Microbiol 2018;3:836–43.

52. Huson DH, Auch AF, Qi J et al. MEGAN analysis of metagenomic data. Genome Res 2007;17:377–86.

53. Kurtz S, Phillippy A, Delcher AL et al. Versatile and open software for comparing large genomes. Genome Biol 2004;5:R12.

54. Manni M, Berkeley MR, Seppey M et al. BUSCO Update: Novel and streamlined workflows along with broader and deeper phylogenetic coverage for scoring of eukaryotic, prokaryotic, and viral genomes. Mol Biol Evol 2021;38:4647–54.

55. Seemann T. Prokka: rapid prokaryotic genome annotation. Bioinformatics 2014;30:2068–9.

56. Emms DM, Kelly S. OrthoFinder: phylogenetic orthology inference for comparative genomics. Genome Biol 2019;20:238.

57. Capella-Gutiérrez S, Silla-Martínez JM, Gabaldón T. trimAl: a tool for automated alignment trimming in large-scale phylogenetic analyses. Bioinformatics 2009;25:1972–3.

58. Zečić A, Dhondt I, Braeckman BP. The nutritional requirements of Caenorhabditis elegans. Genes Nutr 2019;14:15.

59. Cantalapiedra CP, Hernández-Plaza A, Letunic I et al. eggNOG-mapper v2: Functional annotation, orthology assignments, and domain prediction at the metagenomic scale. Mol Biol Evol 2021;38:5825–9.

60. Kanehisa M, Sato Y, Kawashima M. KEGG mapping tools for uncovering hidden features in biological data. Protein Sci 2022;31:47–53.

61. Kanehisa M, Sato Y, Kawashima M et al. KEGG as a reference resource for gene and protein annotation. Nucleic Acids Res 2016;44:D457–62.

62. Jones P, Binns D, Chang H-Y et al. InterProScan 5: genome-scale protein function classification. Bioinformatics 2014;30:1236–40.

63. Jumper J, Evans R, Pritzel A et al. Highly accurate protein structure prediction with AlphaFold. Nature 2021;596:583–9.

64. Wang J, Youkharibache P, Zhang D et al. iCn3D, a web-based 3D viewer for sharing 1D/2D/3D representations of biomolecular structures. Bioinformatics 2020;36:131–5.

65. Weinert LA, Araujo-Jnr EV, Ahmed MZ et al. The incidence of bacterial endosymbionts in terrestrial arthropods. Proc R Soc B Biol Sci 2015;282:20150249.

66. Foster J, Ganatra M, Kamal I et al. The Wolbachia genome of Brugia malayi: Endosymbiont evolution within a human pathogenic nematode. PLOS Biol 2005;3:e121.

67. Haegeman A, Vanholme B, Jacob J et al. An endosymbiotic bacterium in a plant-parasitic nematode: Member of a new Wolbachia supergroup. Int J Parasitol 2009;39:1045–54.

68. Cordaux R, Pichon S, Hatira HBA et al. Widespread Wolbachia infection in terrestrial isopods and other crustaceans. ZooKeys 2012:123–31.

69. Holterman M, Schratzberger M, Helder J. Nematodes as evolutionary commuters between marine, freshwater and terrestrial habitats. Biol J Linn Soc 2019;128:756–67.

70. Jensen P. Feeding ecology of free-living aquatic nematodes. Mar Ecol Prog Ser 1987;35:187–96.

71. Clec’h WL, Chevalier FD, Genty L et al. Cannibalism and predation as paths for horizontal passage of Wolbachia between terrestrial isopods. PLOS ONE 2013;8:e60232.

72. Tian J, Liu J, Zhao H et al. Molecular surveillance reveals a potential hotspot of tick-borne disease in Yakeshi City, Inner Mongolia. BMC Microbiol 2023;23:359.

73. Aivelo T, Norberg A, Tschirren B. Bacterial microbiota composition of Ixodes ricinus ticks: the role of environmental variation, tick characteristics and microbial interactions. PeerJ 2019;7:e8217.

74. Namina A, Kazarina A, Lazovska M et al. Comparative microbiome analysis of three epidemiologically important tick species in Latvia. Microorganisms 2023;11:1970.

75. Daveu R, Laurence C, Bouju-Albert A et al. Symbiont dynamics during the blood meal of Ixodes ricinus nymphs differ according to their sex. Ticks Tick-Borne Dis 2021;12:101707.

76. Engelstädter J, Telschow A. Cytoplasmic incompatibility and host population structure. Heredity 2009;103:196–207.

77. Vandekerckhove T, Watteyne S, Bonne W et al. Evolutionary trends in feminization and intersexuality in woodlice (Crustacea, Isopoda) infected with Wolbachia pipientis (alpha-Proteobacteria). Belg J Zool 2003;133:61–9.

78. Kageyama D, Nishimura G, Hoshizaki S et al. Feminizing Wolbachia in an insect, Ostrinia furnacalis (Lepidoptera: Crambidae). Heredity 2002;88:444–9.

79. Negri I, Pellecchia M, Mazzoglio P j et al. Feminizing Wolbachia in Zyginidia pullula (Insecta, Hemiptera), a leafhopper with an XX/X0 sex-determination system. Proc R Soc B Biol Sci 2006;273:2409–16.

80. Arai H, Anbutsu H, Nishikawa Y et al. Combined actions of bacterio-phage-encoded genes in Wolbachia-induced male lethality. iScience 2023;26, DOI: 10.1016/j.isci.2023.106842.

81. Perlmutter JI, Bordenstein SR, Unckless RL et al. The phage gene wmk is a candidate for male killing by a bacterial endosymbiont. PLOS Pathog 2019;15:e1007936.

82. Kiefer JST, Schmidt G, Krüsemer R et al. Wolbachia causes cytoplasmic incompatibility but not male-killing in a grain pest beetle. Mol Ecol 2022;31:6570–87.

83. Moriyama M, Nikoh N, Hosokawa T et al. Riboflavin provisioning underlies Wolbachia’s fitness contribution to its insect host. mBio 2015;6:10.1128/mbio.01732-15.

